# Compensatory Mechanisms in Visual Sequence Learning: An fMRI Study of Children with Developmental Language Disorder

**DOI:** 10.1101/2025.06.16.659875

**Authors:** Martyna Bryłka, Jakub Wojciechowski, Tomasz Wolak, Hanna B. Cygan

## Abstract

Symptoms of developmental language disorder (DLD) may in part result from an underlying deficit in statistical learning (SL). This learning deficit may be related to the ability to extract probabilistic properties of events in the environment, which is based on the functions of cortical and subcortical brain regions underlying SL. Using a behavioral SL task and functional magnetic resonance imaging (fMRI), we tested SL ability in the visual domain and its neural correlates in children with DLD and their typically developing (TD) peers. During fMRI, children performed SL tasks involving sequences of two types of stimuli: easy-to-name (EN) objects and difficult-to-name (DN) objects. The children underwent a pre-training fMRI, one week of behavioural training and a post-training fMRI. Similar task performance was observed in both groups during the experimental sessions, with an improvement in performance following training in the SL tasks involving both EN and DN objects. FMRI results revealed that, after training, the DLD group presented greater involvement of the frontal cortex and temporal pole for EN objects. Furthermore, in the TD group, the left putamen, globus pallidus (GP) and thalamus were involved in the early stages of SL, whereas in the DLD group, these areas were involved in SL after training. For DN objects, after training, the DLD group presented greater involvement of the parietal and precuneus regions in the SL task performance. Our results suggest that children with DLD may employ different cognitive processes in SL than TD children, possibly as a compensatory mechanism.

## Background

Developmental language disorder (**DLD**) (previously known as specific language impairment [SLI]) is a neurodevelopmental disorder characterized by both quantitative and qualitative language impairments in language production and/or comprehension. Importantly, the language deficits associated with DLD occur in the absence of general cognitive delay. Clinical diagnosis requires additional assessment of nonverbal IQ and the exclusion of other conditions such as autism spectrum disorder, neurological disorders, and hearing impairment (APA, 2013; Leonard, 2014; Bishop et al., 2017). The estimated prevalence of DLD is 5–7% (Norbury et al., 2016). Despite a growing body of research on the symptoms and determinants of the disorder, the precise mechanisms underlying DLD remain unclear. In their review, Abbott and Love (2023) emphasize the considerable heterogeneity of DLD and the limited number of studies investigating its neural basis, which has resulted in conflicting findings.

Scientific literature suggests that linguistic impairments observed in DLD may, at least in part, be related to deficits in statistical learning (**SL**) (Bogaerts et al., 2021; Ullman and Pullman, 2015; Tallerman et al., 2009; Tomblin et al., 2007; Ullman and Pierpont, 2005). In this context, linguistic information can be viewed as statistical events that trigger experience-based learning of phonology, word structure, grammar, and syntax. SL occurs implicitly, based on the probability of particular phonemes, words and syntactic elements appearing in the child’s experience (Kuhl, 2004; Ramscar, 2010). This type of learning begins early in development, and its basic nature has also been demonstrated in neurophysiological animal studies (Schapiro and Turk-Browne, 2015). Sequential SL is considered an essential mechanism for language acquisition, helping individuals recognize language patterns necessary for effective comprehension and production of language (Kuhl, 2004; Ramscar, 2010). It is proposed as a domain-general mechanism, independent of processing modality. SL dysfunction would manifest in DLD as an impairment in language development.

Deficits in SL processes among children with DLD have been documented in behavioral experiments using both verbal and nonverbal stimulus streams (Lum et al., 2014; Obeid, 2016; Lammertink et al., 2017). Lammertink et al. (2020a) demonstrated, using reaction time (**RT**) measures in an online target detection task, that children with DLD are less effective at learning acoustic non-adjacent dependencies compared to typically developing (**TD**) peers. Furthermore, it was shown that word learning through cross-situational learning, which involves tracking co-occurrences between words and their referents, is particularly difficult for children with DLD. These children require greater exposure to learn word-referent pairs compared to TD peers, suggesting an SL deficit that affects lexical-semantic knowledge (Broedelet et al., 2023; Ahufinger et al., 2021). A meta-analysis of studies using the serial RT (**SRT**) paradigm (Lum et al., 2014), which taps into visuoperceptual-motor procedural learning, revealed a significant but small effect size (Standardized Mean Difference = 0.328). Although part of the experiments included in the meta-analysis reported non-significant effects, the joint analysis suggests that DLD is associated with impairments in procedural learning as measured by the SRT task (Lum et al., 2014).

While there is a general consensus that SL deficits contribute to the emergence of DLD symptoms, the exact impact and precise characteristics of these deficits need further verification. For example, even when SL deficits are not evident behaviorally in simple task performance (Lammertink et al., 2020b; Lum et al., 2014), they should be reflected in functional impairments in neuronal networks, including the frontal lobe and basal ganglia (Ullman and Pullman, 2015) that underlie SL related to language acquisition (Tagarelli et al.; 2019). The procedural deficit hypothesis (**PDH**) proposes that impaired procedural memory contributes to DLD (Ullman and Pierpont, 2005; Ullman and Pullman, 2015; Ahufinger et al., 2021). In this context, SL may be considered one of the basic processes underlying procedural learning (Nemeth et al., 2013; Simor et al., 2019). PDH suggested that individuals with DLD suffer from deficits in corticostriatal brain networks underlying procedural learning, with abnormalities in both linguistic and non-linguistic functions. The hypothesis also suggests that variation in language abilities among children with DLD depends on compensatory mechanisms involving declarative learning. Specifically, if the procedural memory system is impaired in most cases of DLD, then language functions would rely more on explicit memorization abilities, which depend on the hippocampus and medial temporal brain structures (Ullman and Pullman, 2015; Lum et al., 2012; Ullman and Pierpont, 2005).

Previous neuroimaging research involving DLD groups has supported the idea of impairments in the anatomy of subcortical structures that may support SL. The largest meta-analysis to date showed the most pronounced effects in the anterior neostriatum (Ullman et al., 2024). Neuroanatomical studies have also shown reduced myelination in the dorsal caudate, which correlates with reduced language performance (Krishnan et al., 2022). However, only a few studies provide data on the functional neural mechanisms of SL in the SLI/DLD population, with neuroimaging experiments primarily focusing on tasks using linguistic material, such as artificial or foreign grammar paradigms, to investigate these mechanisms. Krishnan et al. (2021) found that while children with DLD showed activation in these regions comparable to TD children during a simple language task, subtle differences in the left IFG and caudate nucleus were observed in a subset of DLD participants with the poorest task performance. Plante et al. (2017) demonstrated in their fMRI experiment that adults with SLI/DLD required more time to acquire words in an unfamiliar language. At the same time, they presented hyperactivation in the left superior temporal gyrus (STG), associated with prelexical speech perception and meaningful speech processing, and in the supramarginal gyrus (SMG), involved in semantic processing and articulatory constraints (Price, 2010), suggesting additional effort required for language learning. Regarding child populations, Zwart et al. (2018) presented data from an experiment using electroencephalography (**EEG**) with a sequential SRT task. Although SLI children showed similar behavioral learning as TD children, the patterns of event related potential **ERP** responses differed between the groups. While TD children showed fronto-central P3 enhancement only in the early phase of SL, the SLI/DLD group showed P3 sensitivity in both early and late phases of SL task performance, possibly reflecting compensatory learning processes, related to enhanced top-down cognitive control (Zwart et al., 2018). Studies on the general population suggest the competing character of the involvement of the bottom-up and top-down cognitive processes in SL. Experimental findings from behavioral (Smalle et al., 2022; Tóth-Fáber et al., 2021; Virag et al., 2015; Nemeth et al., 2013) and neuroscientific (Bryłka et al., 2025; Park et al., 2022; Ambrus et al., 2020) studies support this notion. We therefore assumed that in children with DLD, the deficient SL skills may be related to an atypical involvement of the elements of cortico-striatal loop with possible enhanced activity of the cognitively salient prefrontal cortex.

Finally, the statistical learning deficit in DLD is considered to be domain-general and, as such, should refer to basic mechanisms underlying cognitive performance in tasks that engage different processing modalities (Tomblin et al., 2007). In line, the impairment in detecting statistical regularities in the stream of temporally changing information was proven not to be specific to the language domain (Obeid et al., 2016; Lammertink et al., 2017). Taking this into account, as well as possible inter-group differences in lower level central auditory processing (Magimairaj et al., 2021), which could affect the discrimination of stimuli in the auditory SL task performed in the noisy MRI environment, we decided to examine SL mechanisms in DLD using stimuli in the visual modality. We used two SL tasks with abstract and verbalisable objects to account for differences in the involvement of neural circuits between different levels of possible stimulus verbalisation. We assumed that sequences involving abstract stimuli would trigger more implicit SL processes, independent of linguistic support. Conversely, sequences involving verbalisable stimuli may include linguistic cognitive processes that differentiate groups more extensively, favouring TD children in SL (Bryłka et al., 2024). To capture the SL mechanism more closely related to language acquisition, the functional magnetic resonance imaging (fMRI) task design ensured that the information was presented in a purely sequential manner, with no varying spatial locations of the stimuli.

Considering the above, we conducted the first fMRI experiment on the neural mechanisms of visual SL of a probabilistic sequence of objects among children with DLD and a control TD group. In the fMRI paradigm, two types of visual objects were presented: easy-to-name (**EN**) animals and difficult-to-name (**DN**) abstract shapes. We analysed performance-related brain activity and its differences between initial exposure to visual sequences and after one week of training to track inter-group differences in neural processes involved in SL Our experiment aimed to test the following hypotheses:

- Compared to TD peers, children with DLD show atypical involvement of striatal nuclei in the performance of SL tasks.
- Compared to TD peers, children with DLD present compensatory involvement of brain structures underlying declarative memory and top-down cognitive control in SL task performance.
- The differences between the DLD and TD groups are more pronounced in the neural processes underlying the SL of sequences involving verbalisable objects than those involving abstract objects.

## Methods

### Participants

In this longitudinal study, we initially recruited 86 children aged 6;6–9;6 [years; months], including 46 typically developing (TD) children and 40 children with developmental language disorder (DLD). Based on the quality of the fMRI data—specifically excessive head motion or a low signal-to-noise ratio (see neuroimaging section for details)—15 children were excluded from all analyses. An additional 8 children were excluded due to insufficient behavioral responses (fewer than 5 valid responses out of 8 trials in the fMRI session). The final sample included 63 children, of whom one had a co-occurring ASD diagnosis, resulting in a final analysis sample of 62 children, including 27 children with DLD (17 males, 10 females) and 35 TD children (16 males, 19 females).

Identification of children with DLD was based on official documentation provided by psychological and pedagogical counseling centers, including a formal diagnosis and a statement of special educational needs. Additionally, participants underwent a set of psychological assessments to evaluate their linguistic and nonverbal functioning. The children’s language abilities were verified using the Language Development Test (LDT; Pol. *Test Rozwoju Językowego*, Smoczyńska et al., 2015), administered by a certified psychologist. Nonverbal intelligence was assessed using the Fluid Reasoning subtest of the Stanford–Binet Intelligence Scales, Fifth Edition (Roid, 2003), which provides a rapid screening measure of general cognitive functioning. Only children with a documented history of language difficulties and a score below 1.5 SD in total language level or in any of the LDT subscales on the day of testing were included in the DLD group. Additionally, based on interviews with parents and the Autism Spectrum Rating Scales, children with symptoms of autism spectrum disorder or hearing impairment were excluded.

All participants were right-handed and had normal or corrected-to-normal vision, which was ensured through the use of specialized glasses tailored for compatibility with magnetic resonance imaging (**MRI**). The participants did not have any contraindications for undergoing MRI scanning.

Ethical approval for the study was obtained from the Bioethics Committee of the Institute of Physiology and Pathology of Hearing in Warsaw, Poland. The study was conducted in accordance with the Declaration of Helsinki, the ethical principles for medical research involving human subjects. The parents of the children gave informed written consent before starting the study procedures.

## Procedure

### fMRI procedure

The focus of the procedure design was to enable psychologically and physically safe participation of young children and successful stimulation of statistical learning cognitive processes without child’s active effort and irrespective of behavioral response. The children participated in two fMRI sessions (pre-training and post-training), separated by a break of 6 to 8 days, during which they performed online SL training. The fMRI sessions aimed to elicit automatic learning of statistical sequences through the observation of visual symbols. The paradigm used was a modification of an experimental procedure previously designed to stimulate learning of a 1-level statistical sequence (Giorgio et al., 2018; Wang et al., 2017), which was visually adapted to examine children in early elementary school age. The paradigm has been tested with typically developing children, representing a range of language functioning levels, to evaluate its level of difficulty, procedural clarity, and training effectiveness. It consisted of a mixed-design task with four task runs. In each run, participants encountered two types of sequences: statistically ordered and fully random.

Two task runs consisted of four easily identifiable objects: a panda, a lion, a pig, and a frog, representing a linguistic task (**EN**). The other two task runs contained four symbols from the Hebrew alphabet, which were abstract to the participants, representing a non-linguistic task (**DN**) (Figure 1A). The statistical blocks were constructed using a 1-back statistical design, where the appearance of stimulus N was determined by the preceding stimulus (N-1). Following stimulus A, there was an 80% probability that stimulus B would occur and a 20% probability that stimulus C would occur; following stimulus B, there was an 80% probability that stimulus C would occur and a 20% probability that stimulus D would occur, and so on.

**Figure 1.**
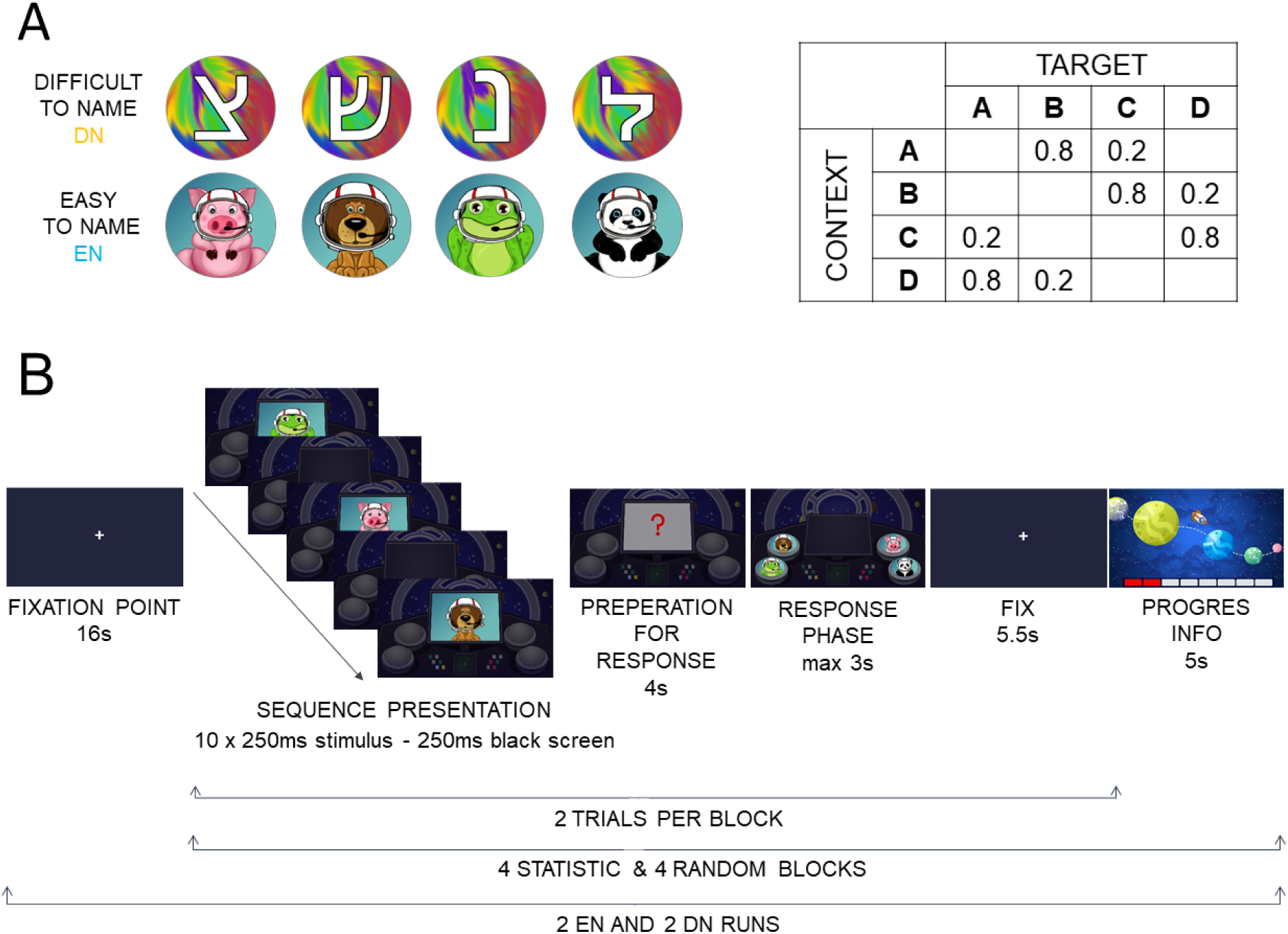
Statistical learning task procedure overview. **(A) The experimental stimuli and their statistical relations in statistical sequences (B) fMRI procedure** During each trial, participants underwent the following sequence of events: Stimulus Presentation Block (Duration: 5.5 seconds): Sequential presentation of visual symbols, either animal pictures or abstract symbols, aimed to provoke unconscious learning of statistical sequences. Each trial featured a sequence of ten stimuli and ten black screens, with each stimulus displayed for 250 ms, followed immediately by a black screen of equal duration before the next stimulus. Pre-response Interval (Duration: 4 seconds): Following the stimulus presentation, a red question mark appeared for one second, indicating that participants should prepare for a response. Response Phase: Participants were instructed to intuitively select the next image in the sequence using a response pad with four buttons within a maximum response time of 3 seconds. Following the response, a fixation point was presented for 5.5 seconds, acting as a short inter-trial interval. Progress Assessment (Duration: 5 seconds): Each block ended with a 5-second period during which participants received visual information on the progress of the task run. This task design, incorporating both easily identifiable objects and abstract symbols, along with deterministic and random sequences, facilitated the investigation of the early stage of implicit learning processes.

**Figure 2.**
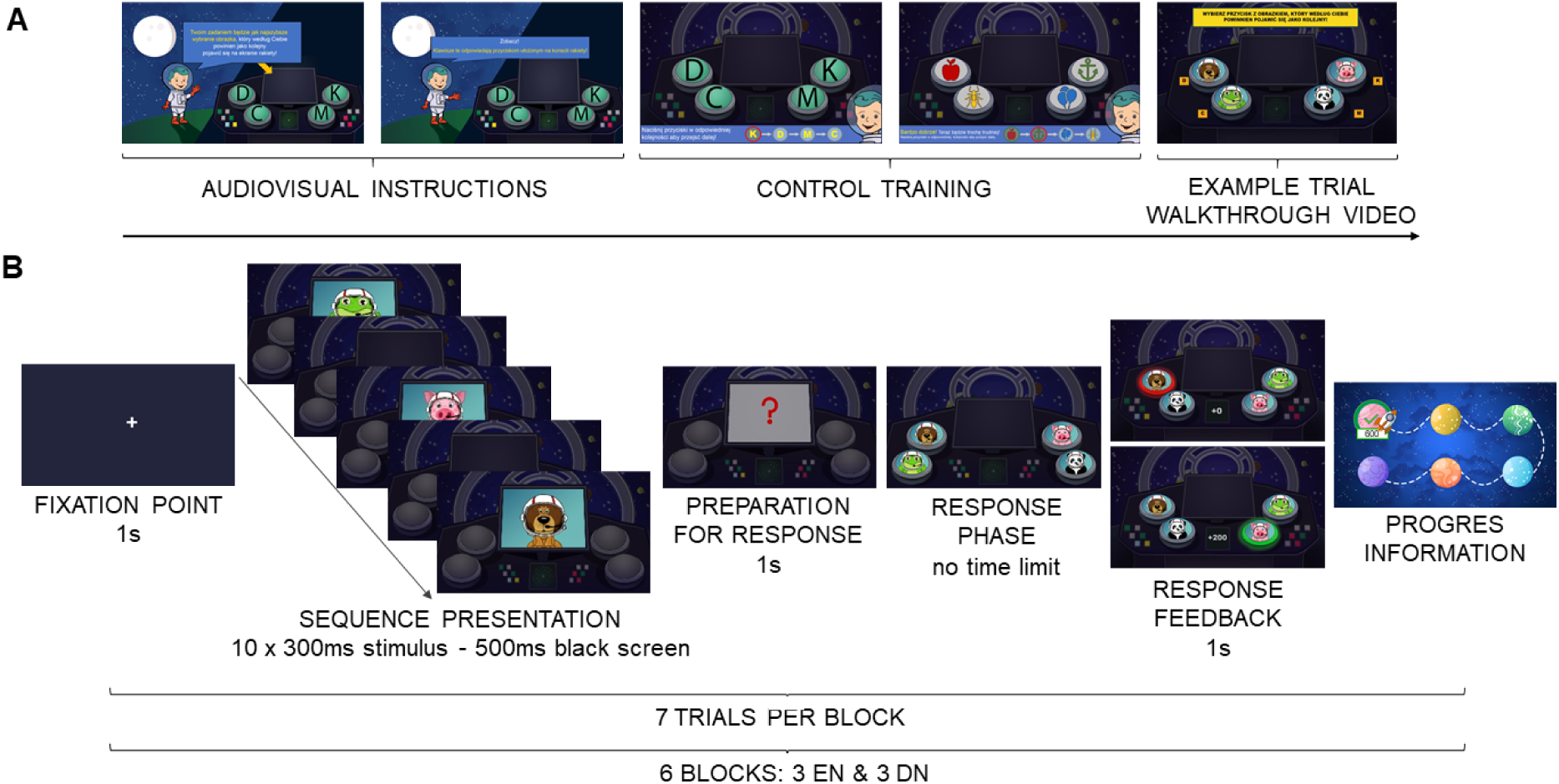
Overview of the online training procedure. **(A)** Visualization of the familiarization phase. Participants first receive **audiovisual instructions** guiding them through the task and explaining the purpose of the training. This is followed by **control training**, which includes a detailed audio-visual introduction to the objectives and flow of the training. In this phase, participants learn how to use the task and familiarise themselves with the interface. Finally, they watch an **example trial walkthrough video**, which illustrates how to complete a trial. ***(B)*** Visualization of the training procedure. Each session begins with a **fixation poin**t displayed for 1,000 ms. This is followed by the **sequence presentation** where stimuli are displayed for 300 ms with a 500 ms black screen in between, repeated 10 times. The sequence is followed by a **preparation for the response phase**, indicated by a red question mark for 1,000 ms, signalling the participant to prepare to respond. In the **response phase**, the participant chooses an answer with no time limit, though visual prompts appear after 2 seconds to encourage quick responses. **Response feedback** is provided for 1,000 ms, displaying whether the selected answer fits the sequence and how many points were earned. The participants can track their **progress** through a progress map at the end of the session. Each block consists of **7 trials**, and each session includes **6 blocks**: 3 with the EN stimulus type and 3 with the DN stimulus type, presented in random order.

Each stimulus appeared on the screen for 250 ms, followed by a black screen for 250 ms before the next stimulus was presented. This design aimed to create a dynamic and probabilistic sequence, fostering a context in which participants could engage in SL. Each task run comprised eight blocks, with four deterministic 1-back statistical blocks and four fully random blocks. Each block consisted of two task trials (Figure 1B).

Prior to the first fMRI session, participants completed a computerized training session consisting of a single task trial involving verbal stimuli presented in a randomized order. The training was conducted using the same response pads as those employed during the actual scanning procedure, in order to familiarize the children with the device and ensure task compliance. During the training session, participants were instructed to observe the rapidly changing stimuli. When the sequence was displayed, and a red question mark appeared, participants were to select one image from a pool of four that they believed should appear next in the sequence. The training continued until the experimenter was sure that the child had fully understood the procedure.

The experimental protocol was controlled by the Psychopy software (Peirce et al., 2019). The stimuli were presented via a mirror mounted to the MR coil and displayed on a full-HD LCD screen (NordicNeuroLab) inside the MRI room. Behavioral responses were collected using MR-compatible response pads (SmitsLab).

### Online training procedure

The participants underwent four online training sessions, structured similarly to the fMRI sessions but without the inclusion of random sequences. Parents were informed about the details of the training procedure and were responsible for initiating the training for their children. Before starting the first home training session, participants completed a control training session. This session included detailed audiovisual instructions explaining the purpose and procedure of the training. Afterward, participants were shown a walkthrough video of an example trial (Figure 3A). Each subsequent training session began with a few practice trials to reinforce the control skills. All instructions during the training were provided multimodally, incorporating infographics, animations, voice instructions, and text. Participants received immediate feedback on their responses. If the chosen answer met the statistical requirements of the sequence (e.g., after Stimulus A, the participant chose Response B or C), the selected button turned green, and the participant earned points (B|A = 200 points, C|A = 100 points). If the response did not meet the statistical expectations of the sequence (e.g., D|A or A|A), the button turned red, and the participant received 0 points. Each training session consisted of six blocks, with children starting either with three blocks of EN stimuli followed by three blocks of DN stimuli, or with three blocks of DN stimuli followed by three blocks of EN stimuli, with the order randomized both between children and across training sessions. Each block contained seven trials, in which a sequence of 10 stimuli appeared, followed by a response selection, where the participant chose the next stimulus that seemed to fit best. Stimuli within the sequence appeared for 300 ms, followed by a 500 ms blank screen. After the sequence, a red question mark was displayed for 1,000 ms to signal the participant that a choice was needed. The RT was unlimited, but if the child did not respond within 2 seconds, the background would begin to gently pulse as a reminder to respond quickly. Feedback was displayed for 1 second after the answer was submitted (Figure 3B).

**Figure 3.**
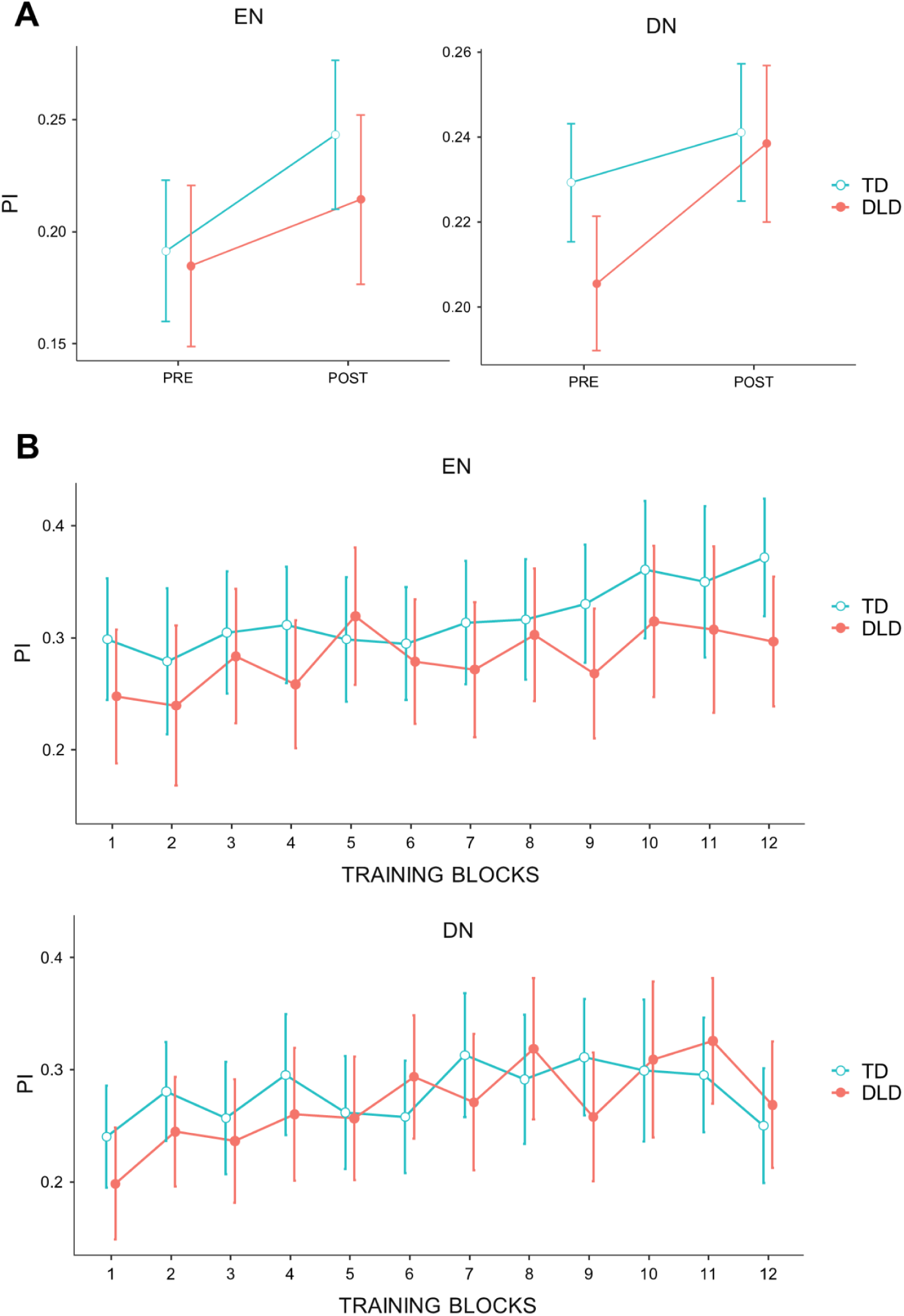
PI Analysis Across fMRI Sessions and Training Blocks. (A) Results of the performance-index (PI) analysis in two tasks: DN and EN, during fMRI sessions, accounting for changes before and after SL training. ANOVA group x time, separately for EN (left panel) and DN (right panel). (B) Changes in PI across 12 training blocks in the groups, separately for EN (top) and DN (bottom).

### Image acquisition

Neuroimaging was performed using a 3 Tesla Siemens Prisma MRI scanner equipped with a 20-channel phased-array RF head coil. Functional data for all tasks were acquired using a multi-band (simultaneous multi-slice) echo-planar imaging sequence (repetition time: TR = 1500 ms, echo time: TE = 27 ms, flip angle = 90°, field of view: FOV = 192 × 192 mm, 94 × 94 mm image matrix, 48 transversal slices of 2.4 mm slice thickness, voxel size of 2.0 × 2.0 × 2.4 mm, slice acceleration factor = 2, in-plane acceleration factor = 2, integrated Parallel Acquisition Technique: iPAT = 4, acquisition time TA = 6:17 min). Structural images were collected using a T1-weighted 3D MP-RAGE sequence (TR = 2300 ms, TE = 2.26 ms, TI = 900 ms, 8° flip angle, FOV = 208 × 230 mm, 232 × 256 mm image matrix, voxel size = 0.9 × 0.9 × 0.9 mm, 208 slices of 0.90 mm slice thickness, TA = 4:53 min).

### Preprocessing

For preprocessing the fMRI data, the Statistical Parametric Mapping 12 (SPM12) package (Wellcome Trust Centre for Neuroimaging, London, UK) and an artifact detection tool (ART; https://www.nitrc.org/projects/artifact_detect) were used. Initially, the DICOM data were converted to NIfTI format (Li et al., 2016). The data then underwent motion correction through realignment, followed by co-registration, spatial tissue segmentation, normalization to the standard MNI anatomical template, and smoothing using a Gaussian filter with a full width at half maximum (FWHM) of 6 mm. After preprocessing, the ART toolbox was further used to verify the number of TRs scrubbed at the time of sequence presentation (statistical or random) and during the fixation point (in relation to contrasts such as Statistic > Fix or Random > Fix). For each run and subject, runs with more than 3 scrubbed TRs out of 8 were excluded from the analysis. Additionally, interpolation was performed, and the temporal signal-to-noise ratio (tSNR) was computed for each run to assess data quality. Only subjects with a mean tSNR above 100 were included in the final analysis, and those with excessively deviating mean or maximum tSNR values were excluded. This conservative threshold reflects the limited scan duration per run (248 TRs) and the use of 2 mm isotropic voxels. According to Murphy et al. (2007), detection of ∼1% BOLD signal changes requires approximately 860 time points at tSNR ∼50, indicating that higher tSNR is critical when fewer volumes are available.

## Data analysis

### Behavioral analysis

To examine training-related effects and potential group differences in statistical learning performance, a repeated measures analysis of variance (RM ANOVA) was conducted on the performance index (PI) as the dependent variable. The model included two within-subject factors: Session (Pre-training vs. Post-training) and Stimulus Type (EN vs. DN), as well as one between-subjects factor: Group (TD vs. DLD). PI quantified the alignment between participants’ response behavior and the statistical structure of the task. On each trial, the smaller value between the expected response probability and the participant’s actual response was treated as a trial score, and these were averaged across trials with weights based on context frequency. Although the PI can theoretically range from 0 to 1, the task design limited both predicted and observed probabilities to discrete values (0, 0.2, 0.8), making 0.8 the maximum attainable score in this paradigm. The methodology for calculating the PI was described in detail by Giorgio et al. (2018). Additionally, to assess changes in learning performance during at-home training, an analogous RM ANOVA was conducted on the PI, with Training Session (12 training blocks) and Stimulus Type (EN vs. DN) as within-subject factors, and Group (TD vs. DLD) as a between-subjects factor. Statistical analyses were performed using Jamovi software 2.3 (2022).

### fMRI analysis

In the first level of analysis, we defined conditions and contrasts for statistical and random sequences, using the passive condition (represented by the fixation point) as a reference for each task run and stimulus type. First-level modeling aggregated data separately for each task run, which included either statistical or random sequences of visual symbols (EN or DN) and pre-response intervals following either sequence type. For motion parameters, we utilized the Friston 24-parameter approach (Friston et al., 1996), which regresses out head motion effects from the realigned data, including six head motion parameters, six head motion parameters from the previous time point, and the 12 corresponding squared terms. Additionally, scrubbing regressors based on a 1.5 mm frame displacement were included to reduce motion-related artifacts, as recommended by the ART toolbox. The canonical hemodynamic response function (**HRF**) without derivatives was applied, and no explicit brain mask was used in the first-level analysis.

For the contrasts of statistical maps, task conditions were aggregated across individual task runs, with equal weights assigned. Because some fMRI sessions were excluded from analysis due to excessive motion or poor-quality behavioral data, the number of task runs varied between subjects. To account for this, we applied appropriate weights to ensure comparisons were made between equivalent numbers of runs. Specifically, subjects with two usable runs were assigned a weight of 1/2, while those with only one usable run were given a weight of 1. To compare sequence types, we created first-level contrasts reflecting the difference between statistical and random sequences (Statistic > Random), separately for EN and DN stimuli. This separation allowed us to identify distinct neural patterns that may be associated with processing stimuli differing in the availability of linguistic support during sequence processing. The resulting contrast images for each participant were then entered into second-level (group) analyses to assess condition-related brain activation across the sample.

### Whole-brain analysis

To examine whole-brain activation patterns associated with statistical learning (SL) and how these patterns are modulated by individual learning performance, we conducted two full-factorial group-level analyses in SPM12—separately for the EN and DN stimulus types—using a 2 (Group: TD vs. DLD) × 2 (Time: Pre-training vs. Post-training) factorial design. Each model included four z-scored covariates representing the individual performance index (PI) for each group and session. The inclusion of these covariates allowed us to test for brain–behavior associations and their modulation by group and training. Analyses were conducted on individual first-level contrast images reflecting the comparison of statistical versus random sequences (Statistic > Random), which directly indexed statistical learning-related activity. To test for neural effects of PI, we computed the F-statistic for the Group × Time interaction on the PI covariate. Additionally, we performed two t-tests comparing the PI covariates of DLD and TD groups at pre- and post-training separately (Pre DLD vs. TD; Post DLD vs. TD). We focused our interpretation on models that included PI covariates, as these allowed us to dissociate neural differences related to underlying learning mechanisms rather than mere performance outcomes. This approach aligns with the goal of identifying altered brain–behavior associations in DLD despite comparable task performance, as suggested by Krishnan et al. (2021). Results from models without PI covariates are reported in the Supplementary Materials. Whole-brain statistical maps were thresholded using a voxel-wise threshold of *p* < .005, combined with a cluster-level family-wise error (FWE) correction at *p* < .05. Pediatric fMRI faces unique challenges such as motion artifacts and non-cooperation, which can affect data quality and the choice of statistical thresholds (Altman and Bernal, 2011). The threshold of *p* < .005 may help mitigate some of these issues by allowing for more lenient detection of true activations. This approach is commonly applied in pediatric fMRI studies (e.g., Lussier and Cantlon, 2017; Schug et al., 2022; Beck et al., 2024). Whole brain activations were visualized with the BrainNet Viewer with the extremum voxel method (Xia et al., 2013) and MRIcroGL (https://www.nitrc.org/projects/mricrogl).

### ROI analysis

To investigate the involvement of striatal structures during the execution of SL tasks in our fMRI paradigm for children, we conducted a region of interest (**ROI**) analysis. Striatal regions are well-documented as being involved in SL (Batterink et al., 2019; Schapiro and Turk-Browne, 2015; Ullman et al., 2024, Bryłka et al., 2025), but their activity may be underestimated in whole-brain statistical comparisons of the blood-oxygen-level-dependent (**BOLD**) response. Structural T1-weighted MRI scans were segmented using the FastSurfer pipeline (Henschel et al., 2020). From these segmentations, individual ROI masks were extracted for the bilateral caudate nucleus, putamen, and globus pallidus. To investigate the internal organization of the caudate nucleus with respect to the temporal dynamics of learning, the caudate was manually divided into two anatomically defined subregions: body and tail (caudal portion), based on previously established boundaries used in adult studies (Bryłka et al., 2025, Choi et al., 2020; Looi et al., 2008). All ROI masks were transformed to MNI space, and average beta values were extracted for each participant and condition from first-level contrast maps generated in SPM12. The contrasts of interest compared the BOLD signal for structured versus unstructured sequences (Statistic > Random), separately for each stimulus type (EN and DN). Mean beta values were computed separately for each ROI (caudate body, caudate tail, putamen, and pallidum), with analyses performed independently for the left and right hemispheres. The ROI activity analyses with PI as a covariate were performed in R v4.3.2 using a generalized linear model (**GLM**) from the standard ‘stats’ package. Only results involving Group—whether as a main effect or in interaction with behavioral PI and/or Time—are reported in the ROI analyses

## Results

### 1. Behavioral results

The analysis of behavioral responses revealed a significant main effect of Time, (*F*(1, 60) = 8.82, *p* = .004, η²□ = .128), indicating improved performance from pre- to post-training with mean increase equal 0.032 (*SE* = 0.01). There was no significant main effect of Group (*F*(1, 60) = 1.22, *p* = .274, η²□ = .020), nor of Stimulus Type (*F*(1, 60) = 2.73, *p* = .104, η²□ = .044), suggesting that the observed improvement was comparable across both participant groups and stimulus conditions. Additionally, none of the interaction effects reached statistical significance: (Time × Group, *F*(1, 60) < 0.001, *p* = .980, η²□ < .001; Stimulus Type × Group, *F*(1, 60) = 0.036, *p* = .851, η²□ = .001; Time × Stimulus Type, *F*(1, 60) = 0.82, *p* = .368, η²□ = .014; and Time × Stimulus Type × Group, *F*(1, 60) = 1.14, *p* = .290, η²□ = .019). These results indicate that all participants, regardless of group or stimulus type, demonstrated a similar pattern of learning over time (see Figure 3, top panel).

To examine how children responded to statistical learning training at home, we analyzed changes in performance across twelve training blocks. The analysis revealed significant main effects of Training Session (*F*(11,561) = 3.01, *p* < .001, η²□ = 0.056), and Stimulus Type (*F*(1,51) = 8.53, *p* = .005, η²□ = 0.143), indicating progressive improvement over time and significantly higher PI values for nameable stimuli (EN) compared to abstract stimuli (DN; *M* difference = 0.027, *t*(51) = 2.92, *p* = .005). No significant effects involving groups were observed (Figure 3, bottom panel). This analysis captured changes in statistical learning performance across twelve blocks of at-home training. Post hoc comparisons showed that performance significantly improved between the early and later training sessions, with the strongest gains from block 1 to blocks 8–11 (e.g., block 8: *t*(51) = –3.12, *p* = .003; block 11: *t*(51) = –3.48, *p* = .001). Differences between adjacent blocks were not significant, suggesting that learning gains accumulated gradually throughout the training.

### 2. fMRI results for EN stimuli (easy-to-name)

Brain–behavior correlations based on PI scores revealed group differences at both the pre- and post-training stages, indicating regions where higher performance was linked to greater neural activation. TD children showed significantly greater activation than children with DLD in the left superior occipital gyrus belonging to the dorsal visual stream (Figure 4, top panel). Following training, children with DLD showed significantly greater PI-related activation than TD peers in a distributed set of regions primarily in the left hemisphere. Significant clusters were observed in the left superior frontal gyrus corresponding to the dorsolateral prefrontal cortex, medial frontal cortex, including ventromedial prefrontal cortex and the frontal pole, anterior cingulate cortex, and anterior orbitofrontal cortex. Additional clusters were identified in the left superior temporal pole, anterior insula, and inferior frontal gyrus pars opercularis (Figure 4, bottom panel).

**Figure 4.**
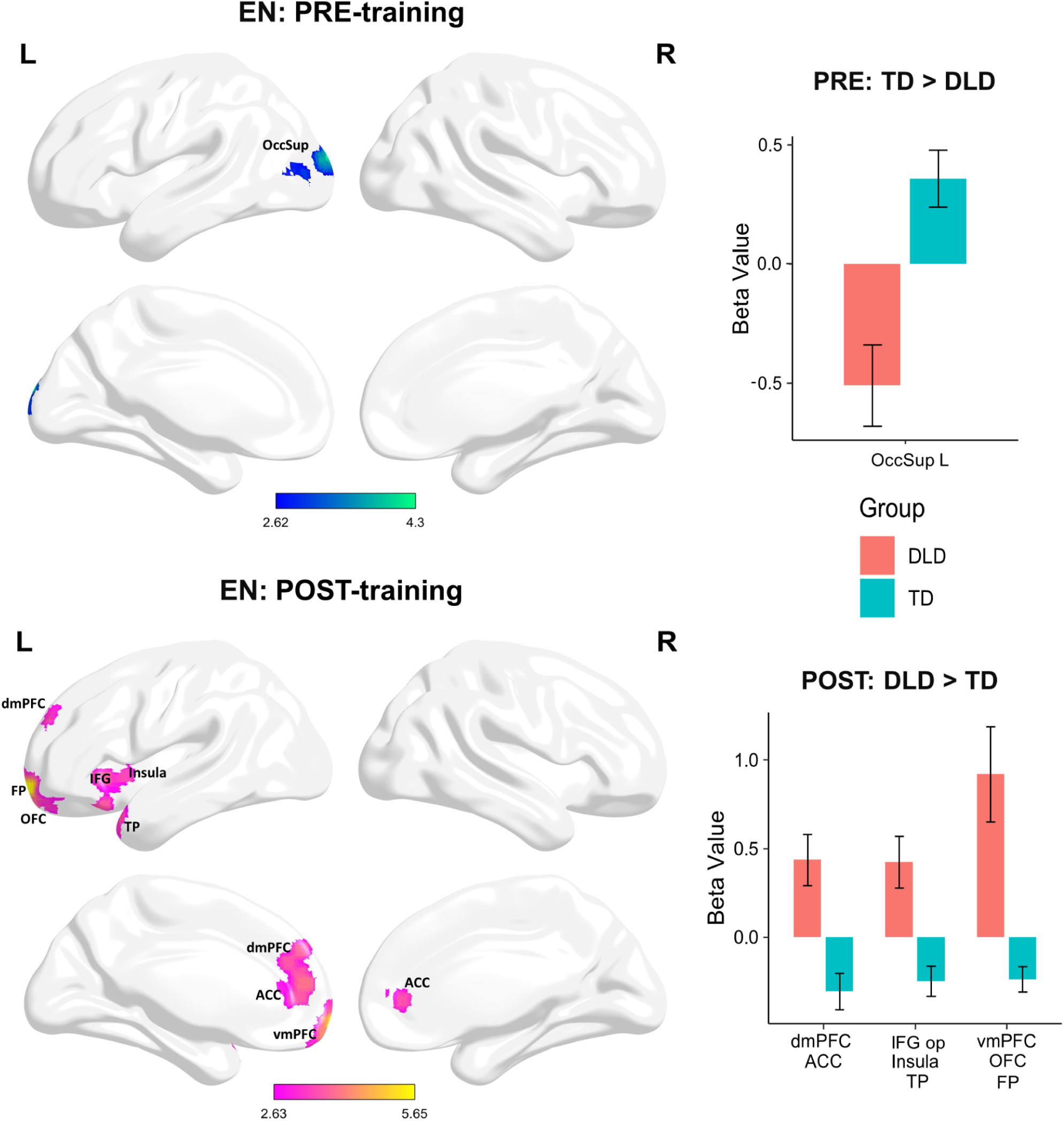
Group differences in PI-modulated activation for EN stimuli before and after training. Both contrasts (TD > DLD and DLD > TD) were tested separately for the pre- and post-training sessions. Panels display only the directions in which significant group differences were observed (p < 0.005, cluster-level FWE-corrected). **Top panel (pre-training):** Left: Surface maps showing clusters with significantly greater PI-related activation in TD compared to DLD participants before training. Right: Mean beta values extracted from the significant cluster identified in the SPM contrast. Values are shown separately for each group (DLD, TD); error bars indicate SEM. **Bottom panel (post-training):** Left: Surface maps showing clusters with significantly greater PI-related activation in DLD compared to TD participants after training. Right: Mean beta values extracted from the significant clusters identified in the SPM contrast. Values are presented separately for each group; error bars represent SEM.

ROI analyses in the EN condition revealed significant group-specific effects in the relationship between behavioral performance and activation only in the left putamen and left pallidum, with training differentially modulating these associations in TD and DLD children. In the left putamen, a significant interaction between PI and group was observed (β = −2.28, SE = 1.10, t(99) = −2.07, p = .041, partial η² = .041), further moderated by time (PI × Group × Time: β = 4.05, SE = 1.54, t(99) = 2.63, p = .010, partial η² = .065). Post-hoc analyses indicated that prior to training, the relationship between PI and activation was moderately positive in TD children (β = 0.64, SE = 0.62), but strongly negative in children with DLD (β = −1.64, SE = 0.91). After training, this pattern reversed: the association became negative in TD (β = −0.91, SE = 0.66) and weakly positive in DLD (β = 0.86, SE = 0.85). Importantly, the DLD group exhibited a statistically significant increase in putamen activation over time (mean difference = 0.39, SE = 0.11, t(99) = 3.55, p = .0006), indicating robust training-related functional changes. These interaction patterns are visualized in Figure 5 (top panel). In the left pallidum, similar effects were found. The PI × Group interaction was significant (β = −2.09, SE = 0.98, t(99) = −2.14, p = .035, partial η² = .044), as was the three-way interaction PI × Group × Time (β = 3.06, SE = 1.36, t(99) = 2.24, p = .027, partial η² = .048). Before training, PI was positively associated with pallidal activation in TD children (β = 0.61, SE = 0.55), but negatively in DLD children (β = −1.47, SE = 0.81). Following training, this reversed: TD children showed a negative association (β = −0.46, SE = 0.58), while the association in DLD children became weakly positive (β = 0.51, SE = 0.75). As in the putamen, a statistically significant increase in activation over time was observed only in the DLD group (mean difference = 0.39, SE = 0.11, t(99) = 2.21, p = .029). Figure 5 (bottom panel) illustrates these group- and time-specific trends.

**Figure 5.**
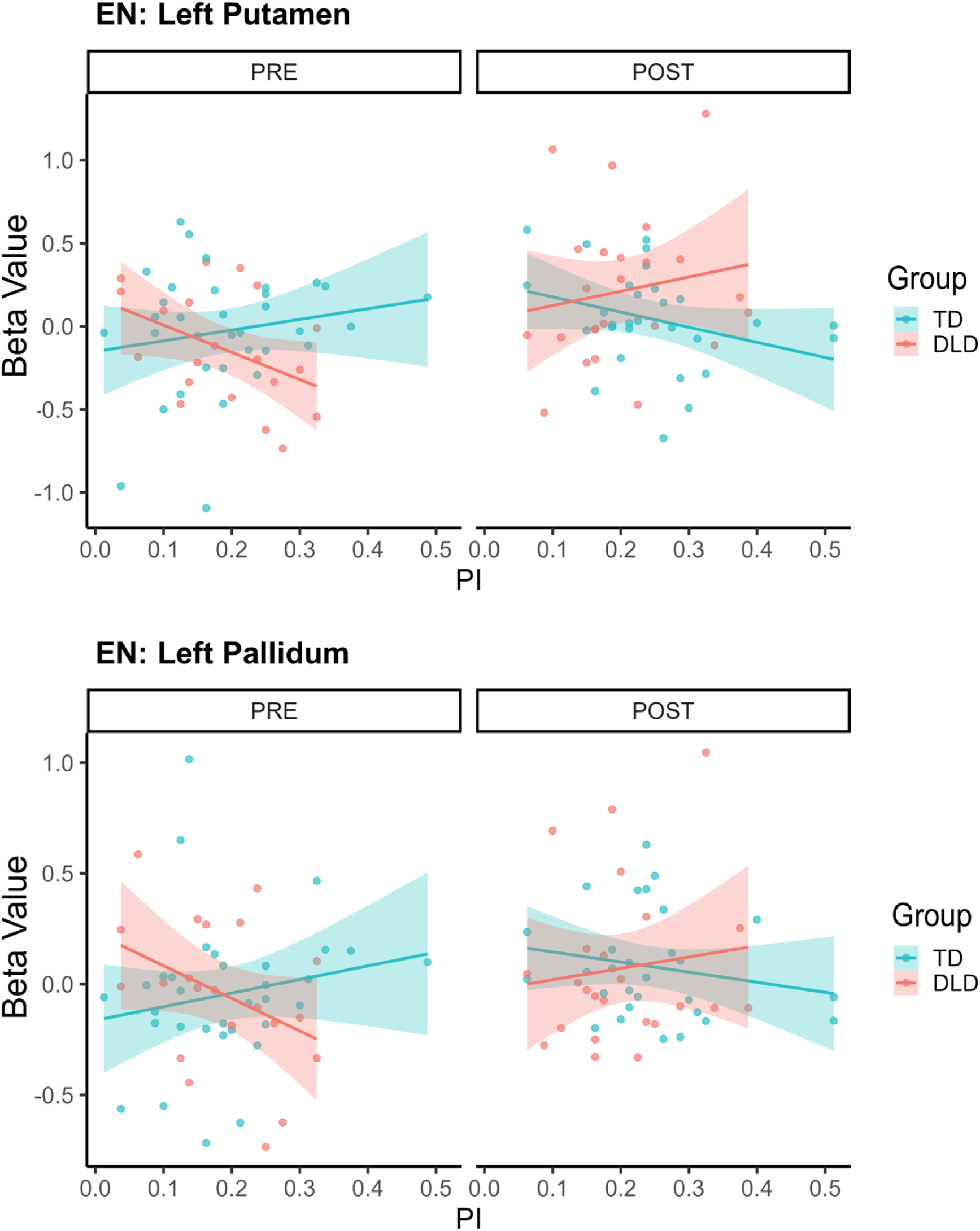
ROI analysis: group-specific associations between PI and neural activation in the left putamen and pallidum (EN stimuli). Scatterplots illustrate the relationship between performance index (PI) and beta values in TD and DLD groups before (PRE) and after (POST) training. Top panel shows data for the left putamen; bottom panel shows data for the left pallidum. Shaded areas represent 95% confidence intervals. The plots correspond to significant PI × Group × Time interactions identified in ROI-based GLM analyses.

### 3. fMRI results for DN stimuli (difficult-to-name)

Performance-based brain–behavior correlations revealed distinct group-level activation patterns in response to DN stimuli. TD children showed significantly greater activation than children with DLD in the anterior ventral portion of the superior frontal gyrus, corresponding to the frontal poles and the medial orbitofrontal cortex located within the ventromedial prefrontal region (Figure 6, top panel). Following training, children with DLD showed significantly greater PI-related activation than TD peers in several parietal regions including the bilateral precuneus and superior parietal lobule (Figure 6, bottom panel).

**Figure 6.**
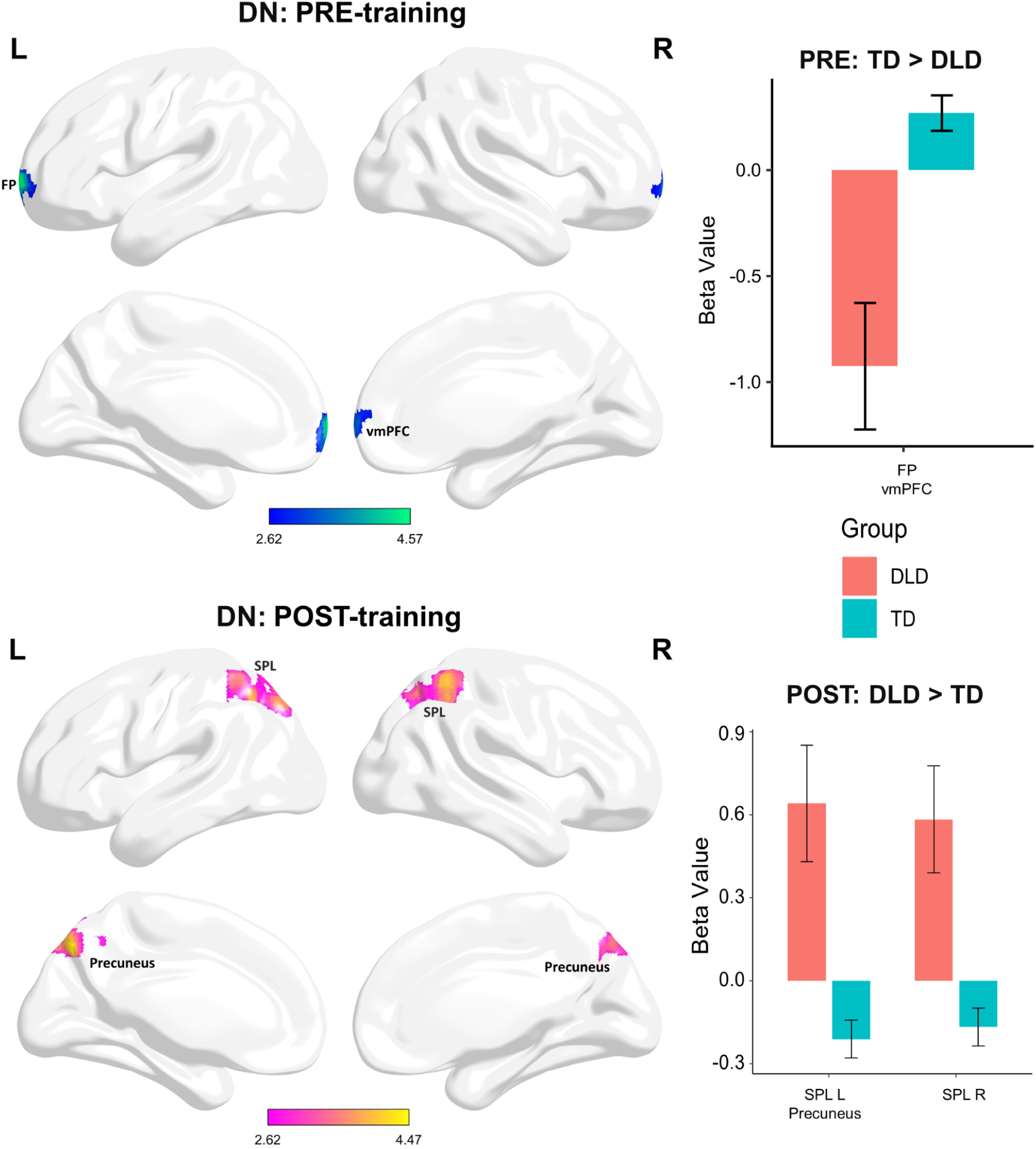
Group differences in PI-modulated activation for DN stimuli before and after training. Both contrasts (TD > DLD and DLD > TD) were tested separately for the pre- and post-training sessions. Panels display only the directions in which significant group differences were observed (p < 0.005, cluster-level FWE-corrected). **Top panel (pre-training):** Left: Surface maps showing clusters with significantly greater PI-related activation in TD compared to DLD participants before training. Right: Mean beta values extracted from the significant cluster identified in the SPM contrast. Values are shown separately for each group (DLD, TD); error bars indicate SEM. **Bottom panel (post-training):** Left: Surface maps showing clusters with significantly greater PI-related activation in DLD compared to TD participants after training. Right: Mean beta values extracted from the significant clusters identified in the SPM contrast. Values are presented separately for each group; error bars represent SEM.

ROI analyses in the DN condition revealed no statistically significant main effects of Group in any of the examined striatal regions. Similarly, no significant PI × Group or PI × Group × Time interactions were found in the caudate body, caudate head, putamen, or pallidum (all p > .10), indicating that striatal responses to statistical input and their relationship to behavioral performance did not differ reliably between TD and DLD groups in this condition.

### 4. Group differences in PI-related brain activation change over time

To examine whether the relationship between individual performance and neural activation was differentially modulated by training across groups, we specifically tested the Group × Time interaction on PI-related activation using a full-factorial model. This approach allowed us to identify brain regions where the association between performance and neural activity changed differently over time in the DLD and TD groups. Although the analysis was designed to detect interaction effects in either direction, significant results indicated stronger training-related increases in PI-dependent activation in the DLD group. For EN stimuli, this interaction followed a distributed pattern involving both subcortical and posterior cortical regions. Greater modulation of performance-related activity over time was observed in the left pallidum, putamen, and mediodorsal thalamus, as well as in posterior visual association areas, including the right calcarine sulcus, middle occipital gyrus, and bilateral middle temporal gyri (Figure 7, top panel). For DN stimuli, a more spatially restricted interaction pattern was observed, again indicating stronger PI-related training effects in the DLD group. Increased activation was found in the left middle occipital gyrus, left inferior parietal lobule, and right precuneus (Figure 7, bottom panel).

**Figure 7.**
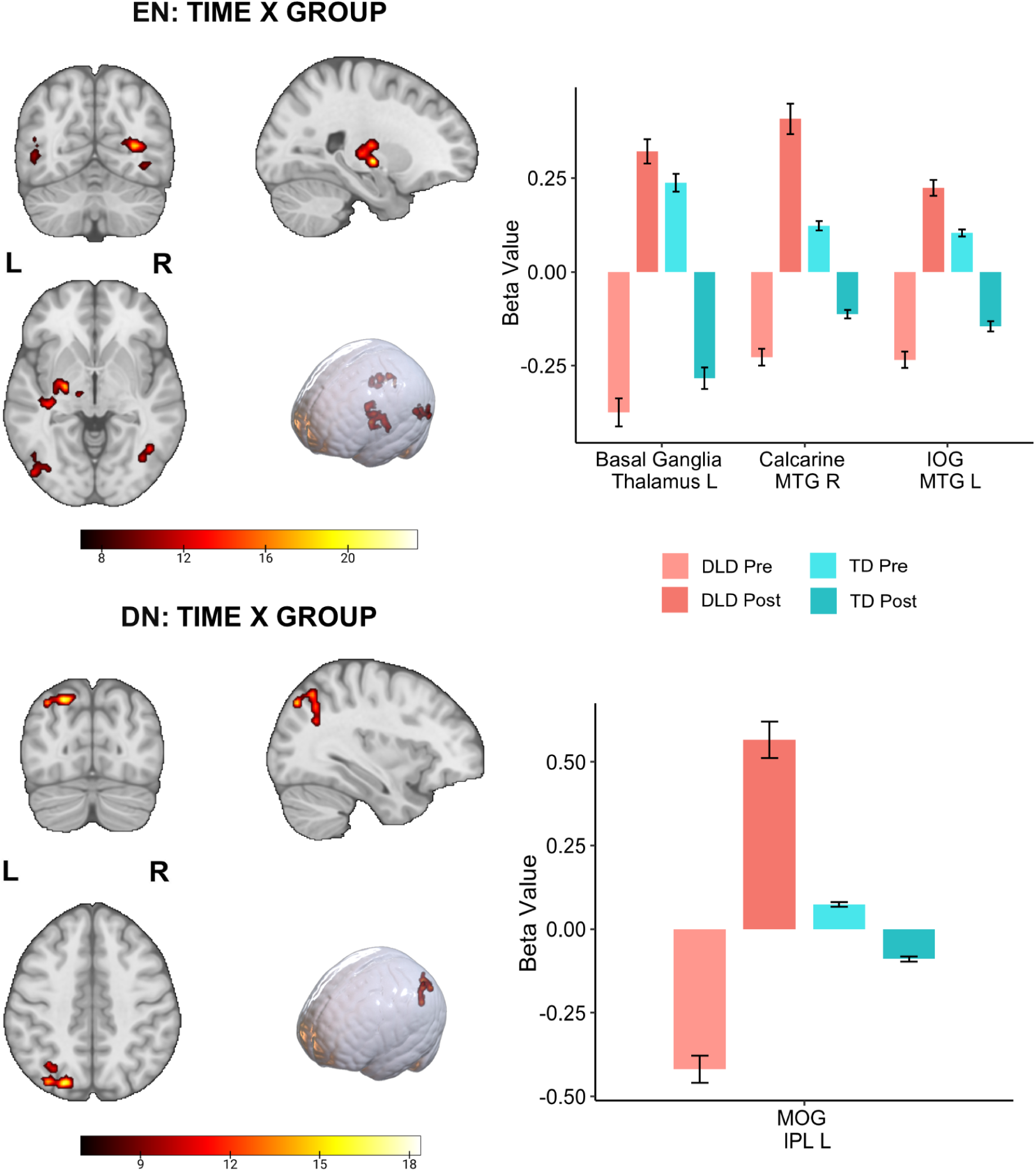
Group × Time interaction in PI-modulated brain activity for EN and DN stimuli. Left: Statistical parametric maps showing brain regions with a significant Group × Time interaction in PI-related activation for easy-to-name (EN; top panel) and difficult-to-name (DN; bottom panel) stimuli. Effects were identified using an F-test testing the interaction between group (DLD vs. TD) and time (pre- vs. post-training) on the PI covariate. Clusters are displayed at p < 0.005 with cluster-level FWE correction. The color scale indicates F-values. Right: Mean beta values representing PI-related activation during pre- and post-training sessions for EN (top panel) and DN (bottom panel) stimuli. Data are shown separately for the DLD and TD groups. Error bars denote SEM.

## Discussion

In the present experiment, we tested SL process in the visual domain and its neural correlates among children with DLD and their TD peers using a behavioral task and fMRI. During the fMRI scanning, children observed transient statistical sequences of two types of stimuli: easy to name (EN) objects and difficult to name (DN) objects, and performed a simple 4-choice SL task. The experimental procedure included a pre-training fMRI session (session 1), one-week behavioral training (4 sessions), and a post-training fMRI session (session 2).

Our findings indicated that children with DLD did not differ from their TD peers in overall behavioral performance. For SL tasks with both types of stimuli EN and DN, analysis of behavioral responses during the pre- and post-fMRI sessions, as well as during home-based training, revealed no significant differences between the groups. Although impairments in SL in DLD are documented in the literature, the lack of effects at the behavioral level has also been observed in other studies (Gabriel et al., 2011; Zwart et al., 2018). A meta-analysis by Lum et al. (2014) found a weak effect, indicating difficulties in SL within the SRT paradigm, which relies on RTs. These inconsistencies in behavioral effects may result from various factors e.g. varying age levels of participants or motor learning difficulties that are more prevalent in DLD than in TD population (Sanjeevan et al., 2015) and thus may impact the performance in SRT tasks. Here, we employed an SL task that did not require RT measurements and featured purely sequential stimulus presentation. It was a modification of the paradigm used in previous fMRI studies of the healthy adult populations (Wang et al., 2017; Giorgio et al., 2018; Bryłka et al., 2025).

Despite similar overall behavioral performance, in the fMRI examination we observed significant differences in brain activity between DLD and TD children related to SL tasks performance. This suggests that the observed fMRI results reflect underlying neural mechanisms involved in SL processing rather than performance-based differences. Krishnan et al. (2021) argued that fMRI paradigms designed to ensure similar performance levels across groups could more effectively capture dysfunctional neural mechanisms in DLD populations. However, in their study, no significant differences were reported in frontostriatal activity during a verb generation task, contradicting the hypothesis of frontostriatal dysfunction in DLD. In contrast, our findings revealed abnormal functions of cortical and subcortical regions in the DLD group. By employing a visual sequential SL paradigm, and by focusing on homogeneous group of young children in their early school years, we were able to identify differences in the brain activity related to task performance in the various cortical regions and basal ganglia, supporting some of the previous expectations (Ullman and Pullman, 2015; Krishnan et al., 2016; Ahufinger et al., 2021).

### Brain mechanisms of SL with EN stimuli

Our investigation focused on differences in SL performance-related brain activity between children with DLD and TD children, and how this activity changes during the learning process. Prior to training, TD children exhibited stronger PI-related activity in the left superior occipital gyrus, suggesting greater visual cortex engagement in the early stage of SL. In contrast, post-training data showed enhanced activation in the DLD group across a left-lateralized network encompassing the superior frontal gyrus corresponding to the dorsolateral prefrontal cortex, medial frontal cortex, including ventromedial prefrontal cortex and the frontal pole, ACC as well as left-lateralized regions involved in language network: IFG, insula and temporal pole. This enhanced frontal and temporal activation during SL of EN objects may indicate involvement of supportive cognitive functions or differences in task-processing strategies employed by children with DLD. Pars opercularis of the left IFG (BA44) is critical for language production and its activation co-occurring with the connected insular areas were associated with the articulatory aspects of speech (Ackermann and Riecker, 2010; Tourville et al., 2019). The DLD group also exhibited post-training increased activation in the left temporal pole (TP), suggesting broader engagement of networks involved in naming and semantic integration (Tsapkini et al., 2011; Campo et al., 2016; Herlin et al., 2021). It is possible that enhanced activity of these structures reflects additional efforts that children with DLD involve during learning sequences with simple objects. They may use inner speech and semantic support (Tourville et al., 2019) in memorisation of visual sequences that are concrete and can be supported by linguistic functions. It should be noted that the pictures used in the SL task could be easily recognised and named by children with DLD. This provided an opportunity for linguistic support that did not exceed their abilities.

Medial frontal cortex and ACC are involved in executive functions including cognitive control, error monitoring and inhibition. They play a pivotal role in encoding congruent events by leveraging stored memory schemas from prior experiences (Guerreiro and Clopath, 2024). They support key cognitive functions such as planning and decision-making, facilitating goal-directed actions (Fuster and Bressler, 2015; Bush et al., 2000; Botvinick, 2007; Heilbronner and Hayden, 2016; Wojciechowski et al., 2024). Specifically, the functions of the medial PFC, include learning associations between input properties (Euston et al., 2012) and predicting action outcomes from learned dependencies (Alexander and Brown, 2011). The involvement of these frontal structures in the DLD group during the post-training session supports the idea that enhanced top-down cognitive control may serve as a compensatory mechanism, enabling children with DLD to achieve similar performance levels to TD children in SL tasks.

At the subcortical level, ROI analyses revealed group-specific associations between SL performance and activity in the left putamen and pallidum. Importantly, while we observed strong effects for those structures, no significant group differences were detected in the caudate nucleus, despite its previously reported role in SL and suggested functional impairments in DLD population (Stillman et al., 2013; Bick et al., 2019; Ullman et al., 2024). This may indicate that learning-related activity in the caudate was similar in both groups of children or was not consistently modulated by performance in our task.

While the TD group showed enhanced activity in the putamen and pallidum during the early stage of SL (pre-training fMRI session), the DLD group recruited these structures only after training, during the second fMRI session. The left putamen has been previously linked to performance in SL tasks with visually presented objects (Hedenius and Persson, 2022; Wang et al., 2017). While the putamen is primarily associated with motor functions and motor sequence learning (Doyon et al., 2009), its involvement in procedural learning highlights its broader role in cognitive processes (Janacsek et al., 2020). Our results may reflect different dynamics and/or learning mechanisms of SL in the studied groups. Delayed activation among children with DLD suggests a shift toward more canonical procedural mechanisms only after extended exposure. Interestingly, putamen activity has also been found to be associated with performance in a sequential visual SL task, particularly following training, in adults (Wang et al., 2017; Giorgio et al., 2018). It is possible that the SL mechanisms in the striatum of children with DLD share similarities with the SL processes exhibited by adults. The literature suggests that implicit SL performance is less efficient in adults than in children under the age of 12 (Janacsek et al., 2012; Nemeth et al., 2013).

The alternative explanation would posit that enhanced late activity of putamen may reflect the additional effort that children with DLD put into their compensatory strategy compared to TD children. For example, it has been shown that multilingual children exhibit increased activation in the left putamen when using a non-native language in which they are not highly proficient. This result was interpreted as reflecting additional effort in controlling articulatory processes in the second language (Abutalebi et al., 2013). The possible compensatory strategy, indicated by enhanced activity in the parts of the left IFG and TP appears to involve semantic processing and object naming. Accompanying putamen activity may reflect additional effort in managing the SL process, which involves top-down control and linguistic/semantic functions.

Further ROI analysis revealed that the pallidum, which is also strongly involved in procedural learning, showed a similar profile of change to the putamen. The pallidum showed enhanced involvement in the TD group in the pre-training session, but contradictory involvement in the DLD group after training. The pallidum is known for its complex connections with the basal ganglia nuclei and thalamus, among other brain regions. Functionally, the pallidum serves as a central hub in the basal ganglia for information processing (Dong et al., 2021), and its specific segments play different roles in behavioral control (Provost et al., 2015). A meta-analysis of fMRI studies using the SRT task for SL examinations in adult populations showed consistent involvement of the pallidum across experiments (Janacsek et al., 2020). Clinical literature additionally indicated pallidum’s role in SL of most implicit probabilistic associations (Sage et al., 2003).

Crucially, the Group × Time analysis of PI-modulated brain activity furtherly confirmed that children with DLD exhibited stronger training-related increases in performance-associated activation than TD children. For EN objects, subcortical effects encompassed not only the left pallidum and putamen but also extended to the medial dorsal thalamus that has multiple connections with prefrontal cortex taking active role in attention orienting, action monitoring and learning (Perry et al., 2022; de Bourbon-Teles et al., 2014; Alexander et al., 1986). Further, Group × Time analysis also revealed enhanced activity in the posterior visual association cortices, including the calcarine sulcus, middle occipital gyrus and bilateral middle temporal gyri, in the DLD group. Thus, the enhanced activity in the pallidum, putamen and thalamus, together with the extensive cortical involvement, appears to indicate a process that becomes overly engaged in children with DLD when they perform successfully in SL, particularly following training. This process may involve more effortful management of multiple cognitive functions, with input information linked to probabilistic sequences.

Previous EEG examination of TD and DLD children during implicit SRT task performance, also suggested enhanced cognitive control among children with DLD. Although no behavioral differences were found, electrophysiological findings suggested that children with DLD (previously SLI) seem to rely on more controlled processes (reflected by a P3 ERP component) in SL, whereas TD children engage both automatic (reflected in the early P2 ERP component) and controlled (late P3 component) processes (Zwart et al., 2018). Similarly, the early fMRI investigation of executive control revealed that children with SLI/DLD recruited frontal and cingulate areas, normally associated with executive control, even when the task did not require them, suggesting compensatory mechanisms for task performance. However, the analysis involved only a small group of 4 children with DLD (Dibbets et al., 2006).

Notably, our study group consisted of well-functioning children with DLD, as evidenced by their nonverbal IQ within the average range (see Table 1). This normative cognitive foundation likely enabled the children with DLD to effectively employ compensatory strategies at the behavioral level.

**Table 1.** Group comparisons and descriptive statistics for language ability, nonverbal IQ, and age.

| Measure | Group | N | Mean | Median | SD | SE | *t* | *df* | *p* | Mean diff. | Cohen's *d* |
| --- | --- | --- | --- | --- | --- | --- | --- | --- | --- | --- | --- |
| LDT (centile) | TD | 35 | 80.23 | 85 | 17.12 | 2.89 | 13.90 | 60 | < .001 | 60.82 | 3.56 |
|  | DLD | 27 | 19.41 | 17 | 17.03 | 3.28 |  |  |  |  |  |
| NV IQ | TD | 35 | 10.74 | 11 | 2.67 | 0.45 | 0.60 | 60 | 0.551 | 0.37 | 0.15 |
|  | DLD | 27 | 10.37 | 10 | 2.06 | 0.4 |  |  |  |  |  |
| Age (years) | TD | 35 | 7.91 | 7.88 | 0.7 | 0.12 | 0.85 | 60 | 0.399 | 0.17 | 0.22 |
|  | DLD | 27 | 7.74 | 7.58 | 0.85 | 0.16 |  |  |  |  |  |

**Note.** LDT = Language Development Test (centile rank); Fluid Reasoning = subscale of the Stanford–Binet Intelligence Scales, Fifth Edition; TD = typically developing; DLD = developmental language disorder. *t*, *df*, and *p* refer to Student's t-test results comparing TD and DLD groups. Cohen's *d* indicates effect size.

**Table 2.**
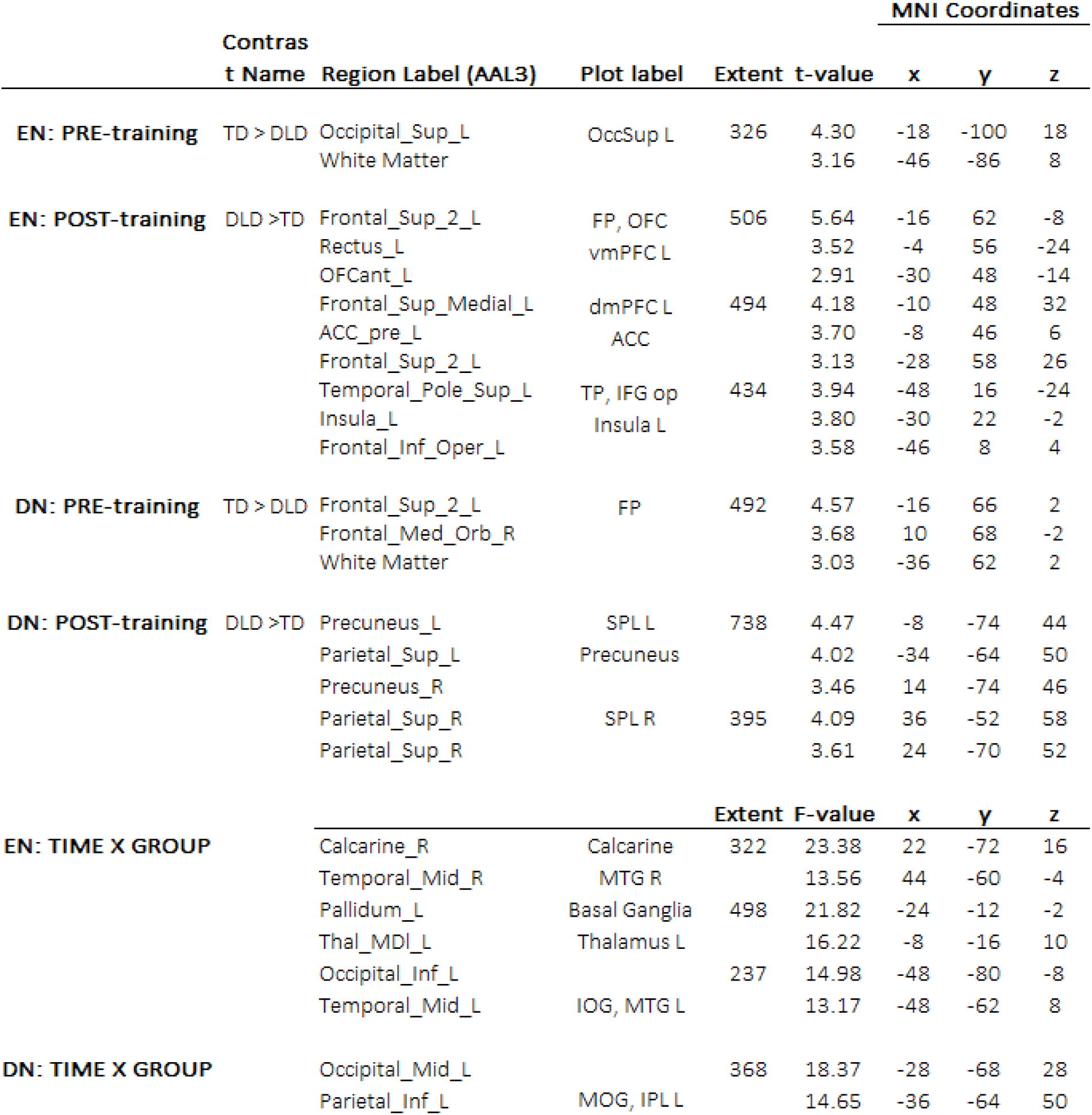
Local Maxima of Group Differences in PI-Related Brain Activation. Table presents brain regions in which activation levels were significantly correlated with the Performance Index (PI) and differed between groups of children with DLD and typically developing (TD) peers. Results are shown separately for DN and EN stimuli across pre- and post-training sessions. Anatomical labels were assigned automatically using the AAL3 atlas. For each contrast, the table lists the relevant brain region(s), abbreviated plot label used for figure visualization, cluster extent (number of voxels), peak t-values, and corresponding MNI coordinates (x, y, z) of the local maxima. The contrasts include: (1) between-group differences at baseline (PRE) and after training (POST); and (2) within-group training-related changes (POST > PRE), with particular focus on group-by-time interactions. Separate analyses were conducted for DN and EN stimuli to differentiate patterns associated with native versus second-language processing. Only clusters exceeding the statistical threshold are reported. The regions are ordered within each contrast by decreasing t-value Note: AAL3 - Automated Anatomical Labelling; EN: Easy-to-name; DN - Difficult-to-name; DLD - Developmental Language Disorder; TD - typically developing; OccSup – Superior Occipital; vmPFC – Ventromedial Prefrontal Cortex; dmPFC – Dorsomedial Prefrontal Cortex; ACC – Anterior Cingulate Cortex; TP – Temporal Pole; IFG – Inferior Frontal Gyrus; SPL – Superior Parietal Lobule; FP – Frontal Pole; OFC – Orbitofrontal Cortex; Insula – Insular Cortex; Precuneus – Precuneus; MTG – Middle Temporal Gyrus; IOG – Inferior Occipital Gyrus; MOG – Middle Occipital Gyrus; IPL – Inferior Parietal Lobule.

### Brain mechanisms of SL with DN stimuli

For sequences with DN stimuli, the fMRI examination of performance-related brain activity revealed that at the early stage of SL, in the pre-training fMRI, there were significant between-group differences in the activity of the bilateral frontal pole. The TD group presented stronger activation of this region related to the effective SL than the DLD group. The PFC plays a pivotal role in encoding congruent events by leveraging stored memory schemas from prior experiences (Guerreiro and Clopath, 2024). FP specifically supports key cognitive functions such as decision-making, and error monitoring, facilitating goal-directed actions (Fuster and Bressler, 2015; Koechlin, 2011). Our results thus show that TD children may be more strongly involved in these cognitive processes at an early stage of SL for effective task performance. Interestingly, as in the EN objects condition, any between-group differences observed during the pre-training session favoured the TD group. However, after training, stronger cortical and subcortical activity was found in children with DLD.

After training, for DN objects in the DLD group we observed bilateral precuneus and superior parietal activity related to task performance. Precuneus activity was found in the subregions previously identified as a part of the para-cingulate network (Dadario and Sughrue, 2023). Recent studies link precuneus with declarative working memory processes including active maintenance and manipulation of information and retrieval of sequential and spatial associations (Flanagin et al., 2023; Dadario and Sughrue, 2023). Additionally, the superior parietal region was associated with sensory integration, attention reorienting, and categorization of abstract objects (Numssen et al., 2021). This further supports the view that children with DLD recruit alternative networks when procedural SL mechanisms are inefficient. Notably, no group differences were observed in striatal activity for DN stimuli, either in whole-brain or ROI analyses suggesting similar levels of their involvement in SL by DLD and TD groups. Alternatively, lack of between-group effects may also be due to the low sensitivity of our measures to subtle and potentially rapid changes in SL-related striatal nucleus activity under the DN condition.

## Summary and conclusions

The most significant findings referred to the SL of sequences with EN objects. We found atypical involvement of subcortical and cortical regions among children with DLD. Examined groups showed an opposite pattern of learning-related change in the activity of the left putamen and pallidum suggesting delayed involvement of basal ganglia possibly reflecting attenuation of implicit aspects of sequential SL among children with DLD. Enhanced involvement of the putamen and the pallidum after SL training in the DLD group was accompanied by extensive fronto-temporal activity, which seems to reflect the enhanced engagement of top-down cognitive control and semantic/linguistic support during the memorisation of concrete visual sequences. However, this compensation mechanism was specific to sequences involving concrete, easily identifiable objects, which enabled support for multiple cognitive processes. For sequences involving abstract DN stimuli, differences in brain activity suggest that children with DLD exhibit fewer supportive mechanisms. The SL was probably driven more implicitly and did not involve extensive top-down cognitive processes. There were also no between-group differences involving subcortical regions during the SL of DN objects, providing no direct evidence of impaired basal ganglia function in children with DLD when processing abstract visual sequences. In the case of DN objects, effects differentiating the DLD group from the TD group were mainly identified in the post-training fMRI, showing performance-related, possibly compensatory, involvement of parietal areas associated with attention and working memory functions.

Taken together, our results on SL processes in children with DLD demonstrate that, despite similar behavioural performance, neurocognitive mechanisms differentiate children with DLD from their TD peers. The neural mechanisms involved in performing a visual SL task with DN and EN objects depend on the level of stimulus abstraction, suggesting that different sets of cognitive functions may be required to support effective SL in children with DLD. Differences between children with DLD and TD are more extensive when SL includes sequences with easily recognisable and nameable objects, leaving open the possibility of compensation through complex cognitive processes involving multiple elements of the cortico-subcortical neuronal circuit. However, future studies are required to improve our understanding of the complex neurocognitive processes involved in SL among children with DLD, and how these processes change across developmental stages.

Our findings may have important implications for clinical practice. They suggest that the performance of children with DLD on implicit SL tasks relevant to language learning is dependent on multiple cognitive functions. The use of alternative cognitive strategies may be effective in overcoming some of the deficits in individual cases. However, the learning demands and individual differences in multiple cognitive abilities should be taken into account.

## Limitations

Our findings must be interpreted in light of certain limitations. First, although the behavioral task design ensured minimal motor demands and emphasized sequential input, its performance metric (PI) may still be influenced by non-learning factors such as attention fluctuations. Second, our DLD sample consisted of relatively high-functioning children with normative nonverbal IQ, which may limit generalizability to the broader DLD population. Third, the use of visual-only stimuli, while controlling for auditory confounds, may not fully capture learning mechanisms engaged during real-world language acquisition. Finally, while the sample size was adequate for fMRI, replication in larger cohorts would strengthen confidence in these results and allow finer subgroup analyses.

## List of abbreviations

SL: statistical learning
DLD: developmental language disorder
TD: typically developing
fMRI: functional magnetic resonance imaging
MRI: magnetic resonance imaging
EN: easy-to-name
DN: difficult-to-name
SLI: specific language impairment
PDH: procedural deficit hypothesis
IFG: inferior frontal gyrus
STG: superior temporal gyrus
EEG: electroencephalography
ERP: event related potential
dmPFC: dorsomedial prefrontal cortex
vmPFC: ventromedial prefrontal cortex
LDT: Language Development Test
tSNR: temporal Signal-to-Noise Ratio
RM ANOVA: repeated measures ANOVA
PI: performance index
REML: restricted maximum likelihood
HRF: hemodynamic response function
FWE: family-wise error
ROI: region of interest
BOLD: blood-oxygen-level-dependent
GLM: generalized linear model
MFG: middle frontal gyrus
MeFG: medial frontal gyrus
ACC: anterior cingulate cortex
MTG: middle temporal gyrus

## Author contributions

All authors contributed to the study conception, design and methodology. Material preparation and data collection were performed by Martyna Byłka and Hanna Cygan. Data analysis was performed by Martyna Bryłka. The first draft of the manuscript was written by Martyna Bryłka and Hanna Cygan and all authors commented on previous versions of the manuscript. All authors read and approved the final manuscript.

## Funding information

This experiment was conducted as a part of the project funded by the Polish National Science Center, grant: UMO-2018/31/D/HS6/03533

## Data availability

Data are available at the following URL: https://identifiers.org/neurovault.collection:19362

## Ethics Approval

Ethical approval for the study was obtained from the Bioethics Committee of the Institute of Physiology and Pathology of Hearing in Warsaw, Poland. The study was conducted in accordance with the Declaration of Helsinki, the ethical principles for medical research involving human subjects.

## Competing interests

Authors declare no financial or non-financial competing interests.

## Notes

https://identifiers.org/neurovault.collection:19362

### Competing Interest Statement

The authors have declared no competing interest.

## References

Abbott, N., Love, T. (2023). Bridging the Divide: Brain and Behavior in Developmental Language Disorder. Brain Sciences, 13(11), 1606. 10.3390/brainsci13111606

Abutalebi, J., Della Rosa, P. A., Gonzaga, A. K. C., Keim, R., Costa, A., Perani, D. (2013). The role of the left putamen in multilingual language production. Brain and language, 125(3), 307–315. 10.1016/j.bandl.2012.03.009

Ackermann, H., Riecker, A. (2010). The contribution (s) of the insula to speech production: a review of the clinical and functional imaging literature. Brain Structure and Function, 214(5), 419–433. 10.1007/s00429-010-0257-x

Ahufinger, N., Guerra, E., Ferinu, L., Andreu, L., Sanz-Torrent, M. (2021). Cross-situational statistical learning in children with developmental language disorder. Language, Cognition and Neuroscience, 36(9), 1180–1200. 10.1080/23273798.2021.1922723

Alexander, G. E., Delong, M. R., Strick, P. L. (1986). Parallel organization of functionally segregated circuits linking basal ganglia and cortex. Annual Review of Neuroscience. 9, 357–381. doi: 10.1146/annurev.ne.09.030186.002041

Alexander, W. H., Brown, J. W. (2011). Medial prefrontal cortex as an action-outcome predictor. Nature neuroscience, 14(10), 1338–1344. 10.1038/nn.2921

Altman, N. R., Bernal, B. (2011). Pediatric applications of fMRI. In Functional Neuroradiology: Principles and Clinical Applications (pp. 545–573). Boston, MA: Springer US. 10.1007/978-1-4419-0345-7_28

Ambrus, G. G., Vékony, T., Janacsek, K., Trimborn, A. B. C., Kovács, G., Nemeth, D. (2020). When less is more: Enhanced statistical learning of non-adjacent dependencies after disruption of bilateral DLPFC. Journal of Memory and Language, 114, 104144. 10.1016/j.jml.2020.104144

American Psychiatric Association, D. S. M. T. F., American Psychiatric Association, D. S. (2013). Diagnostic and statistical manual of mental disorders: DSM-5 (Vol. 5, No. 5). Washington, DC: American psychiatric association.

Beck, J., Chyl, K., Dębska, A., Łuniewska, M., van Atteveldt, N., Jednoróg, K. (2024). Letter–speech sound integration in typical reading development during the first years of formal education. Child Development. 10.1111/cdev.14080

Bick, S. K., Patel, S. R., Katnani, H. A., Peled, N., Widge, A., Cash, S. S., Eskandar, E. N. (2019). Caudate stimulation enhances learning. Brain, 142(10), 2930–2937. 10.1093/brain/awz254

Bishop, D. V., Snowling, M. J., Thompson, P. A., Greenhalgh, T., Catalise-2 Consortium, Adams, C., … House, A. (2017). Phase 2 of CATALISE: A multinational and multidisciplinary Delphi consensus study of problems with language development: Terminology. Journal of child psychology and psychiatry, 58(10), 1068–1080. 10.1111/jcpp.12721

Bogaerts, L., Siegelman, N., Frost, R. (2021). Statistical learning and language impairments: Toward more precise theoretical accounts. Perspectives on Psychological science, 16(2), 319–337. 10.1177/1745691620953082

Botvinick, M. M. (2007). Conflict monitoring and decision making: Reconciling two perspectives on anterior cingulate function. *Cognitive, Affective*, & Behavioral Neuroscience, 7(4), 356–366. 10.3758/CABN.7.4.356

Broedelet, I., Boersma, P., Rispens, J. (2023). Implicit cross-situational word learning in children with and without developmental language disorder and its relation to lexical-semantic knowledge. Frontiers in Communication, 8, 1021654. 10.3389/fcomm.2023.1021654

Bryłka, M., Wojciechowski, J., Wolak, T., Cygan, H. B. (2025). Frontal Deactivation and the Efficacy of Statistical Learning: Neural Mechanisms Accompanying Exposure to Visual Statistical Sequences. Journal of Cognitive Neuroscience, 1–20. 10.1162/jocn_a_02283

Bryłka, M., & Cygan, H. B. (2024). Selective short-term memory impairment for verbalizable visual objects in children with Developmental Language Disorder. Research in Developmental Disabilities, 144, 104637. 10.1016/j.ridd.2023.104637

Bush, G., Luu, P., & Posner, M. I. (2000). Cognitive and emotional influences in anterior cingulate cortex. Trends in cognitive sciences, 4(6), 215–222. 10.1016/s1364-6613(00)01483-2

Campo, P., Poch, C., Toledano, R., Igoa, J. M., Belinchón, M., García-Morales, I., & Gil-Nagel, A. (2016). Visual object naming in patients with small lesions centered at the left temporopolar region. Brain Structure and Function, 221, 473–485. 10.1007/s00429-014-0919-1

Choi, Y., Shin, E. Y., & Kim, S. (2020). Spatiotemporal dissociation of fMRI activity in the caudate nucleus underlies human de novo motor skill learning. Proceedings of the National Academy of Sciences, 117(38), 23886–23897. 10.1073/pnas.2003963117

Dadario, N. B., & Sughrue, M. E. (2023). The functional role of the precuneus. Brain, 146(9), 3598–3607. 10.1093/brain/awad181

de Bourbon-Teles, J., Bentley, P., Koshino, S., Shah, K., Dutta, A., Malhotra, P., … & Soto, D. (2014). Thalamic control of human attention driven by memory and learning. Current biology, 24(9), 993–999. 10.1016/j.cub.2014.03.024

Dibbets, P., Bakker, K., & Jolles, J. (2006). Functional MRI of Task Switching in Children with Specific Language Impairment (SLI). Neurocase, 12(1), 71–79. 10.1080/13554790500507032

Dong, J., Hawes, S., Wu, J., Le, W., & Cai, H. (2021). Connectivity and functionality of the globus pallidus externa under normal conditions and Parkinson’s disease. Frontiers in neural circuits, 15, 645287. 10.3389/fncir.2021.645287

Doyon, J., Bellec, P., Amsel, R., Penhune, V., Monchi, O., Carrier, J., Lehéricy, S., & Benali, H. (2009). Contributions of the basal ganglia and functionally related brain structures to motor learning. Behavioural brain research, 199(1), 61–75. 10.1016/j.bbr.2008.11.012

Euston, D. R., Gruber, A. J., & McNaughton, B. L. (2012). The role of medial prefrontal cortex in memory and decision making. Neuron, 76(6), 1057–1070. doi: 10.1016/j.neuron.2012.12.002. PMID: 23259943; PMCID: PMC3562704.

Flanagin, V. L., Klinkowski, S., Brodt, S., Graetsch, M., Roselli, C., Glasauer, S., & Gais, S. (2023). The precuneus as a central node in declarative memory retrieval. Cerebral Cortex, 33(10), 5981–5990. 10.1093/cercor/bhac476

Fuster, J. M., & Bressler, S. L. (2015). Past makes future: role of PFC in prediction. Journal of cognitive neuroscience, 27(4), 639–654. 10.1162/jocn_a_00746

Gabriel, A., Maillart, C., Guillaume, M., Stefaniak, N., & Meulemans, T. (2011). Exploration of serial structure procedural learning in children with language impairment. Journal of the International Neuropsychological Society, 17(2), 336–343. doi:10.1017/S1355617710001724

Giorgio, J., Karlaftis, V. M., Wang, R., Shen, Y., Tino, P., Welchman, A., & Kourtzi, Z. (2018). Functional brain networks for learning predictive statistics. Cortex, 107, 204–219. 10.1016/j.cortex.2017.08.014

Guerreiro, I. C., & Clopath, C. (2024). Memory’s gatekeeper: The role of PFC in the encoding of congruent events. Proceedings of the National Academy of Sciences, 121(30), e2403648121. 10.1073/pnas.2403648121

Hedenius, M., & Persson, J. (2022). Neural correlates of sequence learning in children with developmental dyslexia. Human Brain Mapping, 43(11), 3559–3576. 10.1002/hbm.25868

Heilbronner, S. R., & Hayden, B. Y. (2016). Dorsal Anterior Cingulate Cortex: A Bottom-Up View. Annual Review of Neuroscience, 39(1), 149–170. 10.1146/annurev-neuro-070815-013952

Henschel, L., Conjeti, S., Estrada, S., Diers, K., Fischl, B. & Reuter, M. (2020). FastSurfer – a fast and accurate deep learning based neuroimaging pipeline. NeuroImage, 219, 117012. doi:10.1016/j.neuroimage.2020.117012

Herlin, B., Navarro, V., & Dupont, S. (2021). The temporal pole: From anatomy to function—A literature appraisal. Journal of chemical neuroanatomy, 113, 101925.

Janacsek, K., Fiser, J., & Nemeth, D. (2012). The best time to acquire new skills: age-related differences in implicit sequence learning across the human lifespan. Developmental science, 15(4), 496–505. 10.1111/j.1467-7687.2012.01150.x

Janacsek, K., Shattuck, K. F., Tagarelli, K. M., Lum, J. A., Turkeltaub, P. E., & Ullman, M. T. (2020). Sequence learning in the human brain: A functional neuroanatomical meta-analysis of serial reaction time studies. NeuroImage, 207, 116387.

Koechlin, E. (2011). Frontal pole function: what is specifically human?. Trends in cognitive sciences, 15(6), 241.

Krishnan, S., Asaridou, S. S., Cler, G. J., Smith, H. J., Willis, H. E., Healy, M. P., & Watkins, K. E. (2021). Functional organisation for verb generation in children with developmental language disorder. NeuroImage, 226, 117599. 10.1016/j.neuroimage.2020.117599

Krishnan, S., Cler, G. J., Smith, H. J., Willis, H. E., Asaridou, S. S., Healy, M. P., & Watkins, K. E. (2022). Quantitative MRI reveals differences in striatal myelin in children with DLD. Elife, 11, e74242. 10.1016/j.neuroimage.2020.117599

Krishnan, S., Watkins, K. E., & Bishop, D. V. (2016). Neurobiological basis of language learning difficulties. Trends in cognitive sciences, 20(9), 701–714. 10.1016/j.neuroimage.2020.117599

Kuhl, P. K. (2004). Early language acquisition: cracking the speech code. Nature reviews neuroscience, 5(11), 831–843. 10.1038/nrn1533

Lammertink, I., Boersma, P., Wijnen, F., & Rispens, J. (2017). Statistical learning in specific language impairment: A meta-analysis. Journal of Speech, Language, and Hearing Research, 60(12), 3474–3486. 10.1044/2017_JSLHR-L-16-0439

Lammertink, I., Boersma, P., Wijnen, F., & Rispens, J. (2020a). Children with developmental language disorder have an auditory verbal statistical learning deficit: Evidence from an online measure. Language Learning, 70(1), 137–178. 10.1111/lang.12373

Lammertink, I., Boersma, P., Wijnen, F., & Rispens, J. (2020b). Statistical Learning in the Visuomotor Domain and Its Relation to Grammatical Proficiency in Children with and without Developmental Language Disorder: A Conceptual Replication and Meta-Analysis. Language Learning and Development, 16(4), 426–450. 10.1080/15475441.2020.1820340

Leonard, L. B. (2014). Children with specific language impairment. MIT press.

Li, X., Morgan, P. S., Ashburner, J., Smith, J., & Rorden, C. (2016). The first step for neuroimaging data analysis: DICOM to NIfTI conversion. Journal of neuroscience methods, 264, 47–56. 10.1016/j.jneumeth.2016.03.001

Looi, J. C., Walterfang, M., Spulber, G., et al. (2008). Volumetrics of the caudate nucleus: Reliability and validity of a new manual tracing protocol. Psychiatry Research: Neuroimaging, 174(1), 67–75.

Lum, J. A., Conti-Ramsden, G., Morgan, A. T., & Ullman, M. T. (2014). Procedural learning deficits in specific language impairment (SLI): A meta-analysis of serial reaction time task performance. Cortex, 51, 1–10. 10.1016/j.cortex.2013.10.011

Lum, J. A., Conti-Ramsden, G., Page, D., & Ullman, M. T. (2012). Working, declarative and procedural memory in specific language impairment. Cortex, 48(9), 1138–1154. 10.1016/j.cortex.2011.06.001

Lussier, C. A., & Cantlon, J. F. (2017). Developmental bias for number words in the intraparietal sulcus. Developmental science, 20(3), 10.1111/desc.12385. 10.1111/desc.12385

Murphy, K., Bodurka, J., & Bandettini, P. A. (2007). How long to scan? The relationship between fMRI temporal signal to noise ratio and necessary scan duration. NeuroImage, 34(2), 565–574. 10.1016/j.neuroimage.2006.09.032

Nemeth, D., Janacsek, K., & Fiser, J. (2013). Age-dependent and coordinated shift in performance between implicit and explicit skill learning. Frontiers in computational neuroscience, 7, 147.

Magimairaj, B. M., Nagaraj, N. K., Champlin, C. A., Thibodeau, L. K., Loeb, D. F., & Gillam, R. B. (2021). Speech perception in noise predicts oral narrative comprehension in children with developmental language disorder. Frontiers in Psychology, 12, 735026. 10.3389/fpsyg.2021.735026

Norbury, C. F., Gooch, D., Wray, C., Baird, G., Charman, T., Simonoff, E., … & Pickles, A. (2016). The impact of nonverbal ability on prevalence and clinical presentation of language disorder: Evidence from a population study. Journal of child psychology and psychiatry, 57(11), 1247–1257. 10.1111/jcpp.12573

Numssen, O., Bzdok, D., & Hartwigsen, G. (2021). Functional specialization within the inferior parietal lobes across cognitive domains. elife, 10, e63591. 10.7554/eLife.63591

Obeid, R., Brooks, P. J., Powers, K. L., Gillespie-Lynch, K., & Lum, J. A. (2016). Statistical learning in specific language impairment and autism spectrum disorder: A meta-analysis. Frontiers in psychology, 7, 1245. 10.3389/fpsyg.2016.01245

Park, J., Janacsek, K., Nemeth, D., & Jeon, H.-A. (2022). Reduced functional connectivity supports statistical learning of temporally distributed regularities. NeuroImage, 260, 119459. 10.1016/j.neuroimage.2022.119459

Perry, B. A. L., Méndez, J. C., & Mitchell, A. S. (2022). Cortico-thalamocortical interactions for learning, memory and decision-making. The Journal of Physiology, 601, 25–35. 10.1113/JP282626

Peirce, J. W., Gray, J. R., Simpson, S., MacAskill, M. R., Höchenberger, R., Sogo, H., Kastman, E., Lindeløv, J. (2019). PsychoPy2: experiments in behavior made easy. Behavior Research Methods. 10.3758/s13428-018-01193-y

Plante, E., Patterson, D., Sandoval, M., Vance, C. J., & Asbjørnsen, A. E. (2017). An fMRI study of implicit language learning in developmental language impairment. NeuroImage: Clinical, 14, 277–285. 10.1016/j.nicl.2017.01.027

Provost, J. S., Hanganu, A., & Monchi, O. (2015). Neuroimaging studies of the striatum in cognition Part I: healthy individuals. Frontiers in systems neuroscience, 9, 140. 10.3389/fnsys.2015.00140

Price, C. J. (2010). The anatomy of language: a review of 100 fMRI studies published in 2009. Annals of the new York Academy of Sciences, 1191(1), 62–88. 10.1111/j.1749-6632.2010.05444.x

Ramscar, M., Yarlett, D., Dye, M., Denny, K., & Thorpe, K. (2010). The effects of feature-label-order and their implications for symbolic learning. Cognitive science, 34(6), 909–957.10.1111/j.1551-6709.2009.01092.x

Roid, G. H. (2003). Stanford–Binet Intelligence Scales, Fifth Edition (SB5). Itasca, IL: Riverside Publishing.

Sage, J. R., Anagnostaras, S. G., Mitchell, S., Bronstein, J. M., De Salles, A., Masterman, D., & Knowlton, B. J. (2003). Analysis of probabilistic classification learning in patients with Parkinson’s disease before and after pallidotomy surgery. Learning & Memory, 10(3), 226–236. http://www.learnmem.org/cgi/doi/10.1101/lm.45903.

Sanjeevan, T., Rosenbaum, D. A., Miller, C., van Hell, J. G., Weiss, D. J., & Mainela-Arnold, E. (2015). Motor issues in specific language impairment: A window into the underlying impairment. Current Developmental Disorders Reports, 2, 228–236. 10.1007/s40474-015-0051-9

Schapiro, A., & Turk-Browne, N. (2015). Statistical learning. Brain mapping, 3(1), 501–506. 10.1016/B978-0-12-397025-1.00276-1

Schug, A. K., Brignoni-Pérez, E., Jamal, N. I., & Eden, G. F. (2022). Gray matter volume differences between early bilinguals and monolinguals: A study of children and adults. Human brain mapping, 43(16), 4817–4834. 10.1002/hbm.26008

Simor, P., Zavecz, Z., Horváth, K., Éltető, N., Török, C., Pesthy, O., Gombos, F., Janacsek, K., & Nemeth, D. (2019). Deconstructing Procedural Memory: Different Learning Trajectories and Consolidation of Sequence and Statistical Learning. Frontiers in psychology, 9, 2708. 10.3389/fpsyg.2018.02708

Smalle, E. H., Daikoku, T., Szmalec, A., Duyck, W., & Möttönen, R. (2022). Unlocking adults’ implicit statistical learning by cognitive depletion. Proceedings of the National Academy of Sciences, 119(2), e2026011119.

Smoczyńska, M., Haman, E., Maryniak, A., Czaplewska, E., Krajewski, G., Banasik, N., … & Morstin, M. (2015). Test rozwoju językowego. Warszawa: Instytut Badań Edukacyjnych.

Stillman, C. M., Gordon, E. M., Simon, J. R., Vaidya, C. J., Howard, D. V., & Howard, J. H. (2013). Caudate Resting Connectivity Predicts Implicit Probabilistic Sequence Learning. Brain Connectivity, 3(6), 601–610. 10.1089/brain.2013.0169

Tagarelli, K. M., Shattuck, K. F., Turkeltaub, P. E., & Ullman, M. T. (2019). Language learning in the adult brain: A neuroanatomical meta-analysis of lexical and grammatical learning. NeuroImage, 193, 178–200. 10.1016/j.neuroimage.2019.02.061

Tallerman, M., Newmeyer, F., Bickerton, D., Bouchard, D., Kaan, E., & Rizzi, L. (2009). Explain and Replicate?. Biological foundations and origin of syntax, 3, 135.

The jamovi project (2022). jamovi. (Version 2.3) [Computer Software]. Retrieved from https://www.jamovi.org.

Tomblin, J. B., Mainela-Arnold, E., & Zhang, X. (2007). Procedural learning in adolescents with and without specific language impairment. Language Learning and Development, 3(4), 269–293. 10.1080/15475440701377477

Tourville, J. A., Nieto-Castañón, A., Heyne, M., & Guenther, F. H. (2019). Functional parcellation of the speech production cortex. Journal of Speech, Language, and Hearing Research, 62(8S), 3055–3070. 10.1044/2019_JSLHR-S-CSMC7-18-0442

Tóth-Fáber, E., Janacsek, K., Szőllősi, Á., Kéri, S., & Nemeth, D. (2021). Regularity detection under stress: Faster extraction of probability-based regularities. Plos one, 16(6), e0253123. 10.1371/journal.pone.0253123

Tsapkini, K., Frangakis, C. E., & Hillis, A. E. (2011). The function of the left anterior temporal pole: evidence from acute stroke and infarct volume. Brain, 134(10), 3094–3105. 10.1093/brain/awr050

Ullman, M. T., & Pierpont, E. I. (2005). Specific language impairment is not specific to language: The procedural deficit hypothesis. Cortex, 41(3), 399–433. 10.1016/S0010-9452(08)70276-4

Ullman, M. T., & Pullman, M. Y. (2015). A compensatory role for declarative memory in neurodevelopmental disorders. Neuroscience & Biobehavioral Reviews, 51, 205–222. 10.1016/j.neubiorev.2015.01.008

Ullman, M. T., Clark, G. M., Pullman, M. Y., Lovelett, J. T., Pierpont, E. I., Jiang, X., & Turkeltaub, P. E. (2024). The neuroanatomy of developmental language disorder: a systematic review and meta-analysis. Nature Human Behaviour, 8(5), 962–975. 10.1038/s41562-024-01843-6

Virag, M., Janacsek, K., Horvath, A., Bujdoso, Z., Fabo, D., & Nemeth, D. (2015). Competition between frontal lobe functions and implicit sequence learning: evidence from the long-term effects of alcohol. Experimental brain research, 233, 2081–2089. 10.1007/s00221-015-4279-8

Wang, R., Shen, Y., Tino, P., Welchman, A. E., & Kourtzi, Z. (2017). Learning Predictive Statistics: Strategies and Brain Mechanisms. The Journal of Neuroscience, 37(35), 8412. 10.1523/JNEUROSCI.0144-17.2017

Wojciechowski, J., Jurewicz, K., Dzianok, P., Antonova, I., Paluch, K., Wolak, T., & Kublik, E. (2024). Common and distinct BOLD correlates of Simon and flanker conflicts which can (not) be reduced to time-on-task effects. Human Brain Mapping, 45(1), e26549. 10.1002/hbm.26549

Xia, M., Wang, J., & He, Y. (2013). BrainNet Viewer: A Network Visualization Tool for Human Brain Connectomics. PLOS ONE, 8(7), e68910. 10.1371/journal.pone.0068910

Zwart, F. S., Vissers, C. T. W., Kessels, R. P., & Maes, J. H. (2018). Implicit learning seems to come naturally for children with autism, but not for children with specific language impairment: Evidence from behavioral and ERP data. Autism Research, 11(7), 1050–1061. 10.1002/aur.1954

