## Supplementary Materials for "Compensatory Mechanisms in Visual Sequence Learning: An fMRI Study of Children with Developmental Language Disorder"

For completeness, we report group-level analyses conducted without the inclusion of PI covariates. These models serve to illustrate overall activation patterns in response to statistical versus random sequences but do not control for inter-individual variability in learning performance. Given the lack of behavioral differences between groups and our primary aim to isolate neural mechanisms underlying statistical learning (SL), we centered our interpretation on covariate-inclusive models (see main text). Two separate 2 (Group: TD vs. DLD) × 2 (Time: Pre-training vs. Post-training) full-factorial analyses were conducted in SPM12, separately for EN and DN stimulus types. First-level contrast images reflecting the comparison between statistical and random sequences (Statistic > Random) were used as input to assess condition-related brain activation across the sample. No behavioral covariates were included in these models. Whole-brain statistical maps were thresholded using a voxel-level threshold of $p < .005$, with cluster-level FWE correction at $p < .05$ to control for multiple comparisons.

### Easy-to-Name stimuli (EN)

The full-factorial model revealed no significant main effect of Group. A significant main effect of Time, independent of Group, was observed in one cluster located in the left angular gyrus (see Table S1). Two clusters showed a significant Group × Time interaction (see Table S2). The first cluster (extent: 282 voxels) was located in the more ventral part of the left prefrontal cortex, including the anterior insula, the orbital part of the inferior frontal gyrus (pars orbitalis), and the middle orbital gyrus. The second cluster (extent: 351 voxels) encompassed more dorsal portions of the left middle frontal gyrus, consistent with the dorsolateral prefrontal cortex (dlPFC). No significant clusters were identified for direct between-group comparisons at either the pre-training or post-training sessions (all $p > .05$, FWE-corrected).

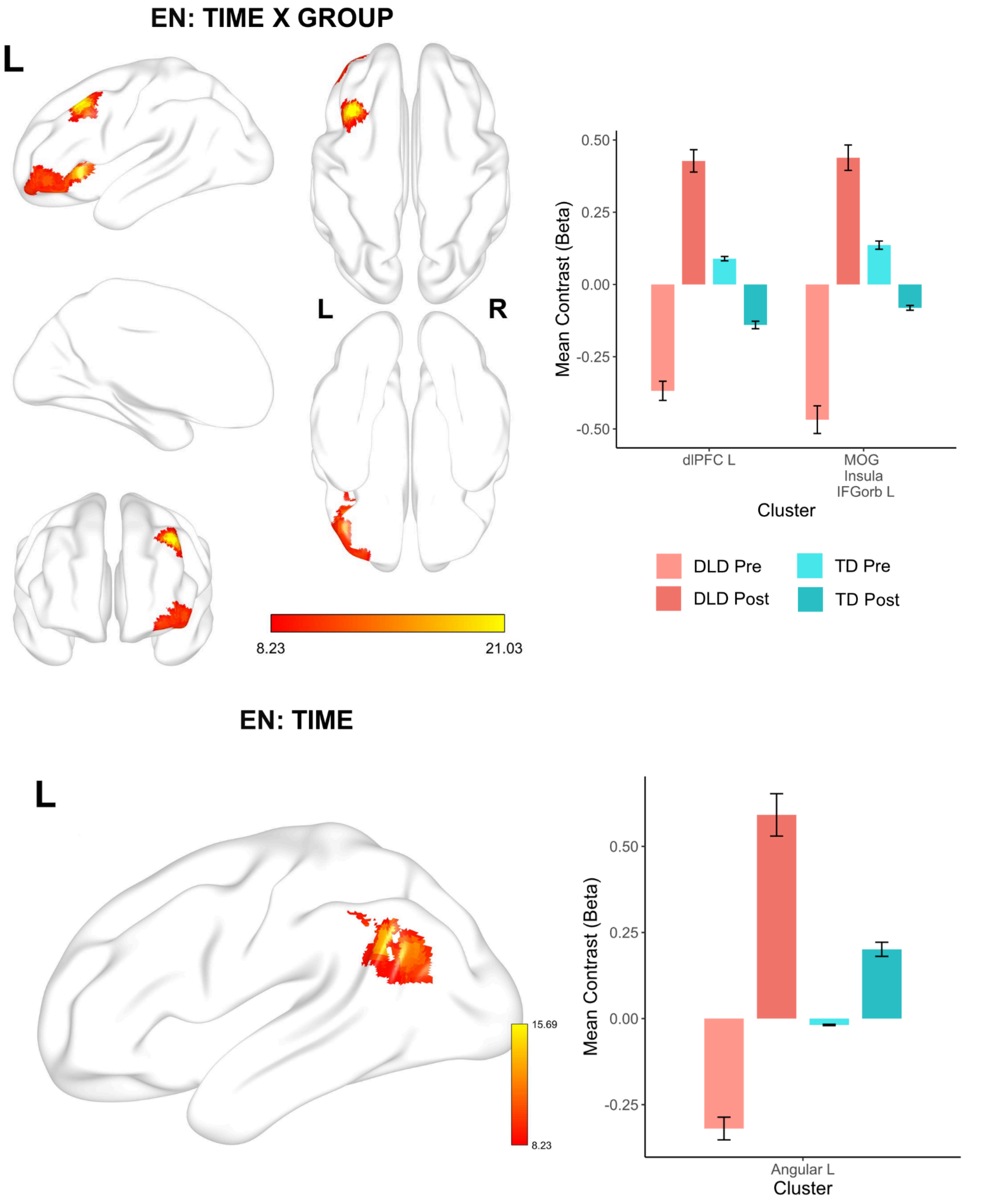


**Figure S1. Main effects and Group × Time interaction in whole-brain activation for EN stimuli (without PI covariate).**

*Top left: Statistical parametric maps showing brain regions with a significant Group × Time interaction in the contrast Statistic > Random for easy-to-name (EN) stimuli. Two significant clusters were identified:*

*one in the more ventral part of the left prefrontal cortex, encompassing the anterior insula, orbital part of the inferior frontal gyrus (pars orbitalis), and middle orbital gyrus; and one in the more dorsal region of the left middle frontal gyrus, consistent with dorsolateral prefrontal cortex (dlPFC).*
*Bottom left: Significant main effect of Time across groups was observed in one cluster located in the left angular gyrus.*
*Bar plots on the right display mean contrast estimates (beta values) for each cluster across pre- and post-training sessions in TD and DLD groups. Statistical maps were thresholded at voxel-level $p < .005$, with cluster-level FWE correction at $p < .05$.*

#### Difficult-to-name stimuli (DN)

No clusters survived FWE correction for the main effects of Group, Time, or the Group × Time interaction. Furthermore, direct comparisons of brain activation between groups for the contrast Statistical > Random (at either pre- or post-training sessions) yielded no significant clusters after FWE correction.

### General effect of the task

To confirm that task performance reliably engaged brain regions involved in sequence processing, we additionally computed contrasts comparing the statistical sequence blocks to baseline fixation (Statistical > Fixation), separately for each stimulus type (EN and DN), averaged across groups and sessions. These contrasts served to illustrate the general effect of task engagement.

It is important to note that both the *Statistical* and *Random* conditions involved the same visual stimuli; however, in the *Statistical* condition, the order of presentation followed a structured transitional probability pattern, while in the *Random* condition, stimuli were presented in a fully randomized sequence. Therefore, the main contrast of interest (Statistical > Random) isolates activation related specifically to the processing of statistical structure, beyond general perceptual or attentional demands. For this reason, only the *Statistical* > *Fixation* contrast is used to visualize the overall effect of the task
All group-level statistical maps and ROI masks are publicly available via **NeuroVault** under the following link: [https://identifiers.org/neurovault.collection:19362](https://identifiers.org/neurovault.collection:19362).

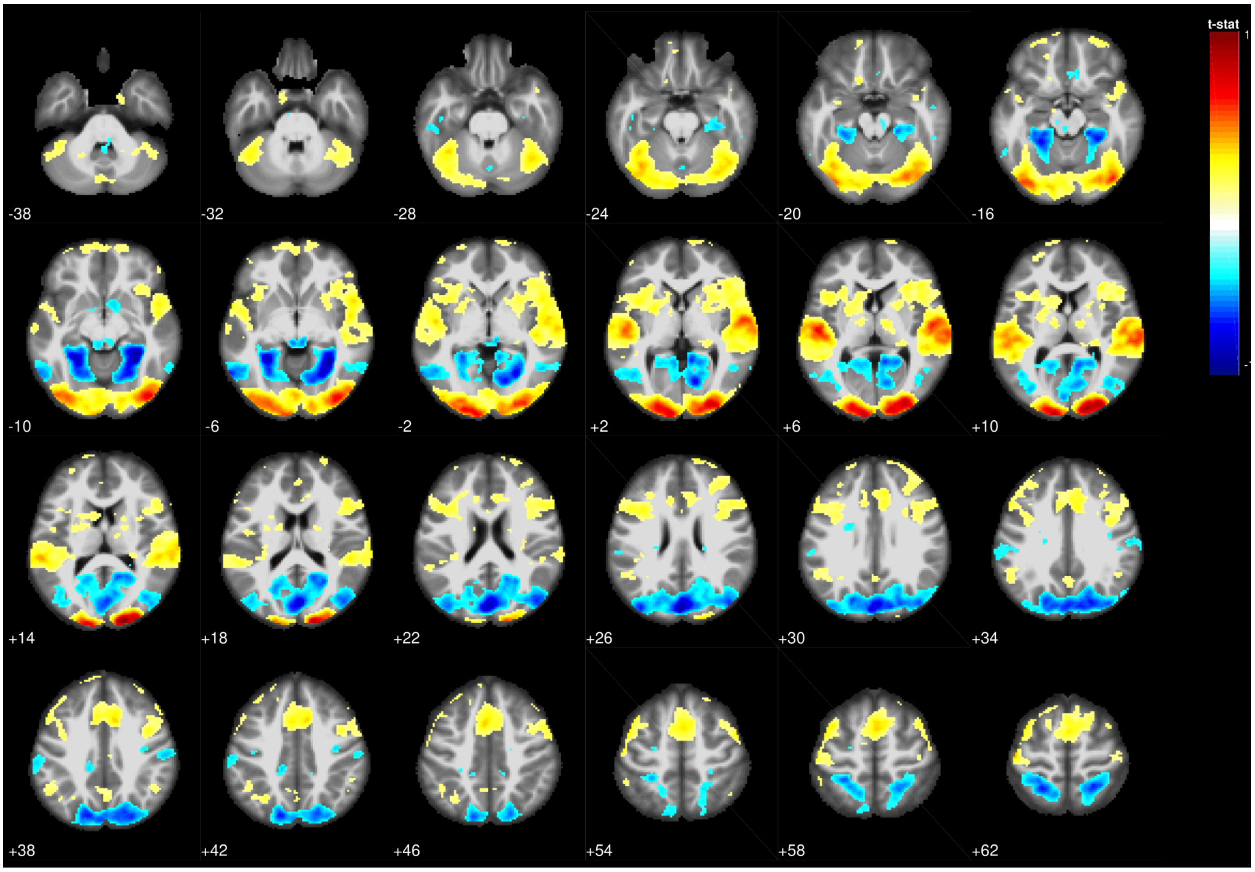


**Figure S2. General effect of task for EN stimuli**

*Statistical parametric maps illustrating the general effect of task engagement for easy-to-name (EN) stimuli. The contrast compared statistical sequence blocks to baseline fixation (Statistical > Fixation), averaged across groups (TD and DLD) and sessions (pre- and post-training). Widespread activation was observed in bilateral occipital cortex, ventral temporal areas, and cerebellum. Additional clusters extended into bilateral parietal and frontal regions. Maps are displayed at a voxel-level threshold of* $p < .005$*, uncorrected for multiple comparisons, and are intended for illustrative purposes only.*

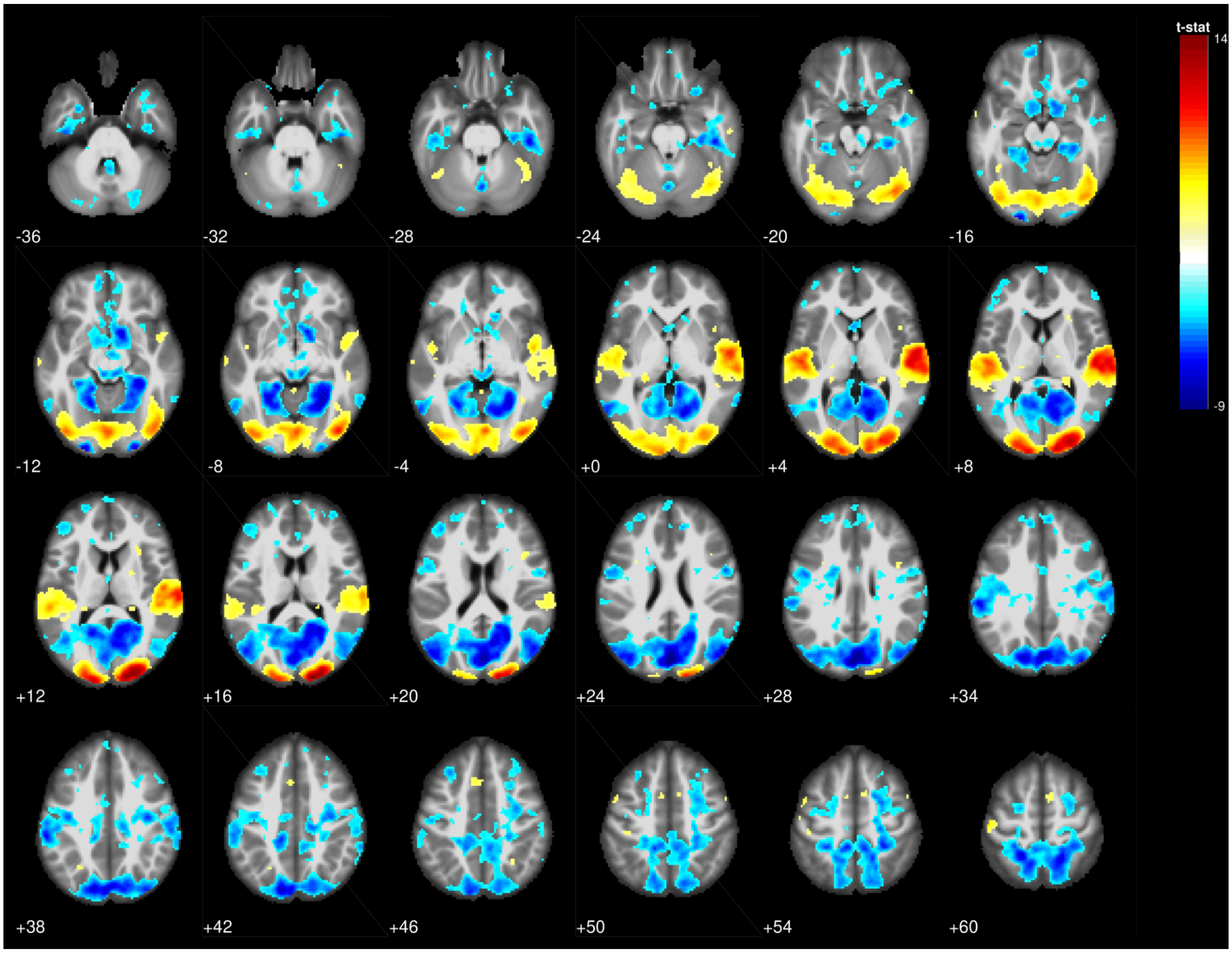


**Figure S3. General effect of task for DN stimuli**

*Statistical parametric maps illustrating the general effect of task engagement for difficult-to-name (DN) stimuli. The contrast compared statistical sequence blocks to baseline fixation (Statistical > Fixation), averaged across groups (TD and DLD) and sessions (pre- and post-training). Widespread activation was observed in bilateral occipital cortex, ventral temporal areas, and cerebellum. Additional clusters extended into bilateral parietal and frontal regions. Maps are displayed at a voxel-level threshold of* $p < .005$*, uncorrected for multiple comparisons, and are intended for illustrative purposes only.*
